# A genome-wide arrayed CRISPR screen identifies PLSCR1 as an intrinsic barrier to SARS-CoV-2 entry that recent virus variants have evolved to resist

**DOI:** 10.1101/2024.02.16.580725

**Authors:** Jérémie Le Pen, Gabrielle Paniccia, Volker Kinast, Marcela Moncada-Velez, Alison W. Ashbrook, Michael Bauer, H.-Heinrich Hoffmann, Ana Pinharanda, Inna Ricardo-Lax, Ansgar F. Stenzel, Edwin A. Rosado-Olivieri, Kenneth H. Dinnon, William C. Doyle, Catherine A. Freije, Seon-Hui Hong, Danyel Lee, Tyler Lewy, Joseph M. Luna, Avery Peace, Carltin Schmidt, William M. Schneider, Roni Winkler, Elaine Z. Yip, Chloe Larson, Timothy McGinn, Miriam-Rose Menezes, Lavoisier Ramos-Espiritu, Priyam Banerjee, John T. Poirier, Francisco J. Sànchez-Rivera, Aurélie Cobat, Qian Zhang, Jean-Laurent Casanova, Thomas S. Carroll, J. Fraser Glickman, Eleftherios Michailidis, Brandon Razooky, Margaret R. MacDonald, Charles M. Rice

## Abstract

Interferons (IFNs) play a crucial role in the regulation and evolution of host-virus interactions. Here, we conducted a genome-wide arrayed CRISPR knockout screen in the presence and absence of IFN to identify human genes that influence SARS-CoV-2 infection. We then performed an integrated analysis of genes interacting with SARS-CoV-2, drawing from a selection of 67 large-scale studies, including our own. We identified 28 genes of high relevance in both human genetic studies of COVID-19 patients and functional genetic screens in cell culture, with many related to the IFN pathway. Among these was the IFN-stimulated gene *PLSCR1*. PLSCR1 did not require IFN induction to restrict SARS-CoV-2 and did not contribute to IFN signaling. Instead, PLSCR1 specifically restricted spike-mediated SARS-CoV-2 entry. The PLSCR1-mediated restriction was alleviated by TMPRSS2 over-expression, suggesting that PLSCR1 primarily restricts the endocytic entry route. In addition, recent SARS-CoV-2 variants have adapted to circumvent the PLSCR1 barrier via currently undetermined mechanisms. Finally, we investigate the functional effects of PLSCR1 variants present in humans and discuss an association between PLSCR1 and severe COVID-19 reported recently.

## Introduction

Viruses maintain a complex relationship with their host cells, co-opting host factors for their replication while being targeted by cellular defense mechanisms. Such cellular defenses include the interferon (IFN) pathway, where the infected cell senses foreign molecules and secretes IFN to trigger an antiviral state in neighboring cells [1].

Approximately 1-5% of critical COVID-19 patients have mutations that compromise the production of or response to type I IFNs, while an additional 15% possess autoantibodies that neutralize type I IFNs [2–8]. This highlights the essential role of type I IFN in the defense against the SARS-CoV-2 virus that caused the COVID-19 pandemic [9, 10]. Consequently, investigating IFN-stimulated genes (ISGs) is crucial to our understanding of the remarkable antiviral systems that evolved in nature and could enhance our preparedness for future pandemics.

Several recent studies have identified ISGs restricting SARS-CoV-2. Most of these studies involved gain-of-function genetic screens, over-expressing individual ISGs. The factors bone marrow stromal cell antigen 2 (BST2), cholesterol 25-hydroxylase (CH25H), lymphocyte antigen 6 family member E (LY6E), 2’-5’-oligoadenylate synthetase 1 (OAS1), and receptor transporter protein 4 (RTP4) were notably identified as SARS-CoV-2 antivirals in these studies [10–15]. One advantage of the gain-of-function approach is that it circumvents potential genetic redundancies between ISGs [16, 17]. However, this approach is biased towards ISGs that act autonomously when over-expressed and does not mimic the cellular context of the IFN response, where hundreds of genes and gene products are differentially regulated to establish an antiviral state. To counter this limitation, two recent publications examined the effects of ISG loss of function in IFN-treated cells. They conducted pooled CRISPR knockout (KO) screens in cells pre-treated with IFN before SARS-CoV-2 infection [18, 19]. By sorting for cells with high SARS-CoV-2 viral load, they identified SARS-CoV-2 restriction factors such as death domain associated protein (DAXX).

Here, we conducted a human whole-genome arrayed CRISPR KO screen to identify genes that influence SARS-CoV-2 infection in cells with or without pretreatment with a low dose of IFN. The arrayed approach, though logistically challenging, has advantages over the pooled format in capturing both proviral and antiviral genes, genes affecting virus egress, and those coding for secreted products that exert their impact on neighboring cells. It reliably captures genotype-phenotype correlations while also unveiling the effects of single gene perturbation on cell growth and death [20]. Additionally, the shorter culture time and lack of competition among cells with different gene KO in the arrayed screen allow the inclusion of genes that would be depleted and deemed essential in a pooled format [21]. The arrayed format thus enables the identification of crucial cellular functions that may be co-opted by the virus or are vital for the cell’s defense against infection.

We then compiled a comprehensive list of genes interacting with SARS-CoV-2, incorporating findings from our own screen as well as existing literature. This meta-analysis revealed several host genes of interest, both previously described and novel. Notably, the ISG product phospholipid scramblase 1 (PLSCR1) emerged as a prominent antiviral factor. PLSCR1 is involved in several biological processes [22], including regulating the movement of phospholipids between the two leaflets of a cell membrane (lipid scrambling) [23] and IFN signaling in the context of virus infection [24].

Follow-up experiments revealed that PLSCR1 is a cell intrinsic factor that restricts spike-mediated SARS-CoV-2 entry, independently of the IFN pathway, via currently undetermined mechanisms. Our genetic screen data and meta-analysis provide a valuable resource to broaden our understanding of coronavirus infection and innate immunity. Furthermore, we extend the recent characterization of PLSCR1 as an antiviral against SARS-CoV-2 impacting COVID-19 outcomes (**Supp Fig. 1**)[19, 25, 26].

## Results

### A genome-wide arrayed CRISPR KO screen identifies known and novel factors influencing SARS-CoV-2 infection

While the liver is not the primary target organ of SARS-CoV-2 infection, human hepatocellular carcinoma Huh-7.5 cells naturally express SARS-CoV-2 dependency factors, including the receptor angiotensin converting enzyme 2 (ACE2), and proved unexpectedly useful in SARS-CoV-2 research [19, 27–34]. Huh-7.5 cells are defective in virus sensing and do not commonly produce IFN during infection [35]. We confirmed that Huh-7.5 cells do not induce ISG expression during SARS-CoV-2 infection (**Fig 1A, Supp Tables 2 and 3**). This is likely due to a defect upstream of IFN production, as these cells did induce ISG expression when treated with recombinant IFN (**Fig 1B, Supp Tables 4 and 5**). Thus, Huh-7.5 cells are a convenient model for studying controlled IFN responses during viral infection. Furthermore, IFN treatment restricted SARS-CoV-2 (**Fig 1C**), indicating that some ISGs are effective in limiting SARS-CoV-2 infection in Huh-7.5 cells. This allows us to study the functional landscape of SARS-CoV-2 restriction, examining both intrinsic factors and those induced in response to IFN signaling. Similar to A549-ACE2 cells [11, 36], Huh-7.5 cells do not express transmembrane serine protease 2 (TMPRSS2) (**Supp Fig 1B**). As a result, SARS-CoV-2 entry is restricted to the endocytic route [37, 38].

**Figure 1.**
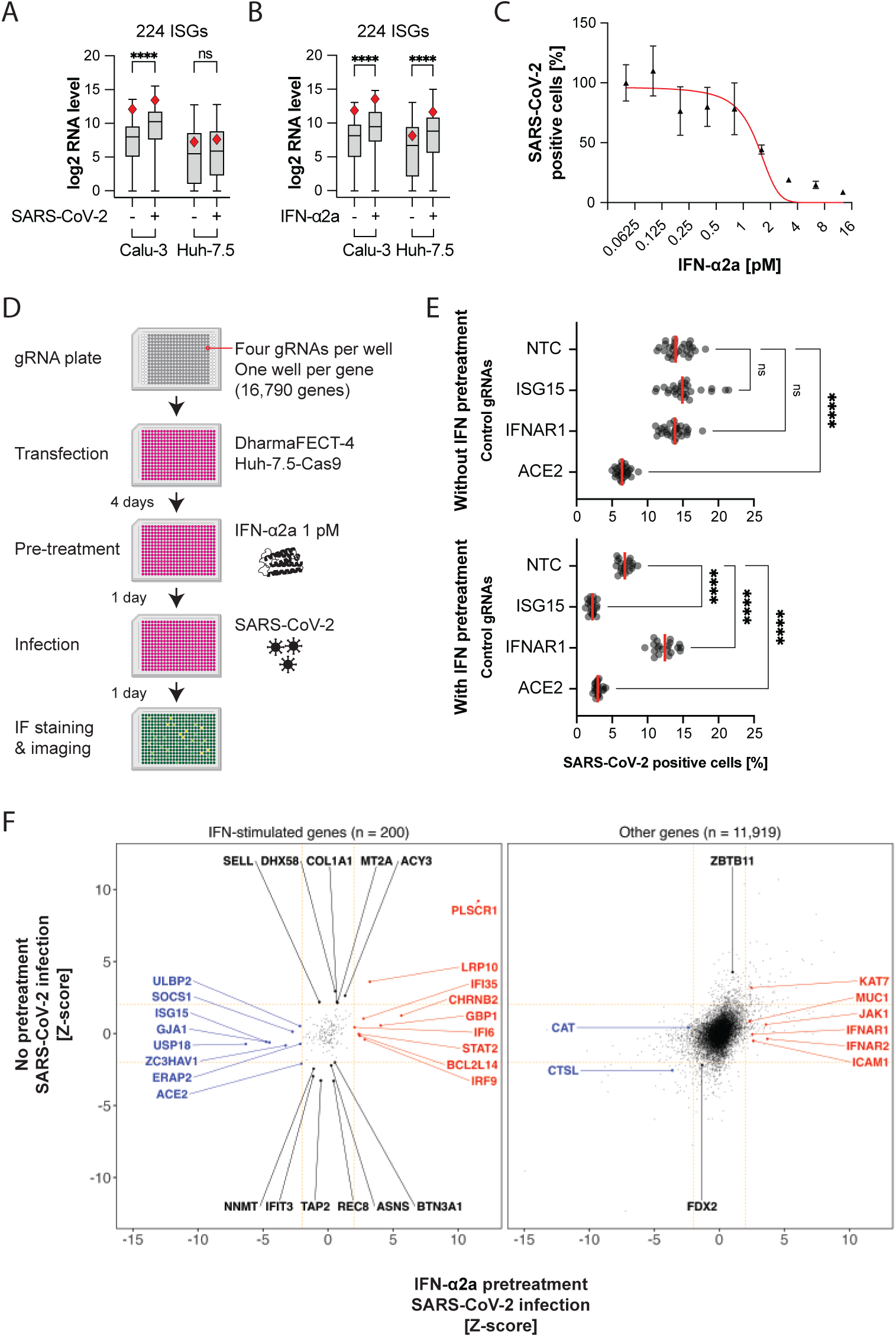
An unbiased arrayed CRISPR KO screen identifies IFN-dependent and IFN-independent genes influencing SARS-CoV-2 infection. A. mRNA-seq comparison between Huh-7.5 and Calu-3 cells, focusing on a subset of 224 ISGs, in response to 24 h SARS-CoV-2 infection MOI 0.03. Red diamond, PLSCR1 RNA level. Viral RNA levels were comparable in both cell lines (not shown). ****, p ≤ 0.0001; two-tailed t-test. B. Cells were treated with 0.5 nM IFN-α2a for 24 h. mRNA-seq analysis as in **(A)**. C. Huh-7.5-Cas9 cells were pretreated with different amounts of IFN-ɑ2a, then infected with SARS-CoV-2 for 24 h followed by IF staining for SARS-CoV-2 N protein; n = 6 separate wells infected on the same day; error bars represent SEM; ****, p ≤ 0.0001; two-tailed t-test. D. Diagram of the arrayed CRISPR KO screen method. E. Median SARS-CoV-2 positive cell percentages determined by IF staining for the indicated control genes. The control genes were distributed across 28 separate wells (without IFN pretreatment) or across 20 separate wells (with IFN pretreatment) for each screened library plate. NTC: non-targeting control. F. The virus level (percentage of infected cells normalized and z-score calculated) is plotted for 24 h 0 pM (y axis) or 1 pM (x axis) IFN-α2a pretreatment followed by 24 h infection (n ≥ 5). The genes were categorized as ISG or other based on mRNA-seq of IFN-α2a-treated cells as in **(B)**. ISGs were defined by a fold change ≥ 2 and padj ≤ 0.05 in the IFN-treatment versus untreated pairwise comparison. The data underlying this Figure can be found in **Supp Table 1**.

Using these cells, we conducted a whole-genome arrayed CRISPR KO screen designed to identify both SARS-CoV-2 proviral and antiviral genes, whose KO reduces or enhances SARS-CoV-2 infection, respectively. In particular, we aimed to identify factors involved in the IFN response, from IFN sensing to ISG induction, including effector ISGs that directly influence the virus life cycle. With this arrayed approach performed in 384-well plates, the cells in each well received a pool of four gRNAs targeting a single host gene. Each well was then either treated with 1 pM IFN-α2a or left untreated before being infected with SARS-CoV-2, followed by SARS-CoV-2 nucleoprotein (N) immunofluorescence staining and high content microscopy (**Fig 1D**). This low dose of IFN was chosen to mimic a cellular environment where IFN triggers an antiviral state before infection. We anticipated that a saturating amount of IFN would lead to high ISG transcription and functional redundancy between effectors, biasing hits towards factors in IFN signaling. In contrast, a low dose of IFN, around the IC50, might enable identification of the specific roles of individual effector ISGs.

We conducted the functional assays four days after KO to allow sufficient time for full protein depletion, considering both cell division and protein half-life (**Supp Fig 2E**).

We included control gRNAs with known proviral and antiviral effects in our screening plates. As expected, the SARS-CoV-2 receptor angiotensin-converting enzyme 2 (ACE2) [9] behaved as a proviral gene with and without IFN pretreatment. The interferon-alpha/beta receptor alpha chain 1 (IFNAR1) [39–41] was essential for the IFN-mediated restriction of SARS-CoV-2. The ISG15 ubiquitin-like modifier (ISG15) has been previously described as a negative regulator of the IFN response. ISG15 stabilizes ubiquitin-specific peptidase 18 (USP18) [42–44], which in turn binds to the interferon-alpha/beta receptor alpha chain 2 (IFNAR2) and blocks signal transduction [45, 46]. Accordingly, ISG15 was proviral in IFN-treated cells (**Fig 1E**).

Of the 16,790 screened genes, we selected 16,178 genes where KO did not lead to changes in cellular fitness, as assessed by nuclei count (−2 ≤ z-score ≤ 2) (**Supp Fig 2B**). Of these, we selected 12,119 genes expressed in three cell lines relevant for SARS-CoV-2 research (A549, Calu-3, Huh-7.5 cells) and human lung cells, the primary target cell type *in vivo*, for downstream analysis [47–49](Supp Table 6). We then binned the genes into two groups for data visualization, depending on whether they were induced by IFN-α2a treatment in Huh-7.5 cells as determined by mRNA-seq (log 2 FC ≥ 2 and padj ≤ 0.05) (**Supp Fig 2C, Supp Tables 7-9**).

We classified 448 genes with a SARS-CoV-2 infection z-score ≥ 2 as antiviral hits and 507 genes with a SARS-CoV-2 infection z-score ≤ −2 as proviral hits (**Supp Table 9**).

Our screen found known and previously unidentified host factors influencing SARS-CoV-2 infection (**Fig 1F**). As expected, positive regulators of IFN signaling, such as IFNAR1,2 [39–41], interferon regulatory factor 9 (IRF9) [50, 51], Janus kinase 1 (JAK1) [52], and signal transducer and activator of transcription 2 (STAT2) [53, 54] were antiviral only in IFN pretreated cells. Known negative regulators of IFN signaling, such as ISG15 [42–44], suppressor of cytokine signaling 1 (SOCS1) [55], and USP18 [45, 46] were proviral only in IFN pretreated cells.

The receptor ACE2 was required for infection [9]. In our mRNA-seq analysis, IFN treatment was found to significantly upregulate ACE2 mRNA levels (**Supp Fig. 2C**). Prior studies indicate that IFN induces transcription of a truncated ACE2 isoform, rather than the full-length receptor for SARS-CoV-2 [56, 57]. The lysosomal cysteine protease cathepsin L (CTSL), required for SARS-CoV-2 spike protein activation during endocytosis [58–60], was also a proviral hit in our screens (**Fig 1F**).

The screen data likely contains false negatives. For example, STAT1 and tyrosine kinase 2 (TYK2) [61, 62] did not influence infection alongside other positive regulators of IFN signaling, which we attribute to the fact that some gRNAs in the library may have not efficiently directed Cas9 to cut at their respective target gene loci.

Collectively, the identification of known proviral and antiviral factors confirms the validity of our screening method.

We performed a Gene Set Enrichment Analysis (GSEA) to identify cellular pathways exhibiting proviral or antiviral properties in our screen. The full GSEA results, including the genes driving each pathway enrichment (so-called *leading edge*), can be found in **Supp Table 10**. Some top pathways ranked by adjusted p-value are summarized in **Supp Fig 2D**. Notably pathways associated with RNA pol II transcription and mRNA maturation, as well as pathways related to cellular respiration, exhibited antiviral activity independent of IFN (**Fig 2**). Surprisingly, RNA pol III transcription, in part driven by the genes RNA polymerase III subunit A (*POLR3A*) and RNA polymerase III subunit B (*POLR3B*), were critical to the antiviral response mediated by IFN. Conversely, factors involved in translation, such as eukaryotic translation initiation factor 3 subunits F and G (EIF3G and EIF3F), likely co-opted for producing viral proteins, were identified as proviral. Similarly, factors regulating cholesterol homeostasis, likely crucial for SARS-CoV-2 entry [14, 63], were also identified as proviral. For instance, the gene sterol regulatory element binding transcription factor 2 (*SREBF2*) was one of the top proviral genes (**Fig 2**).

**Figure 2.**
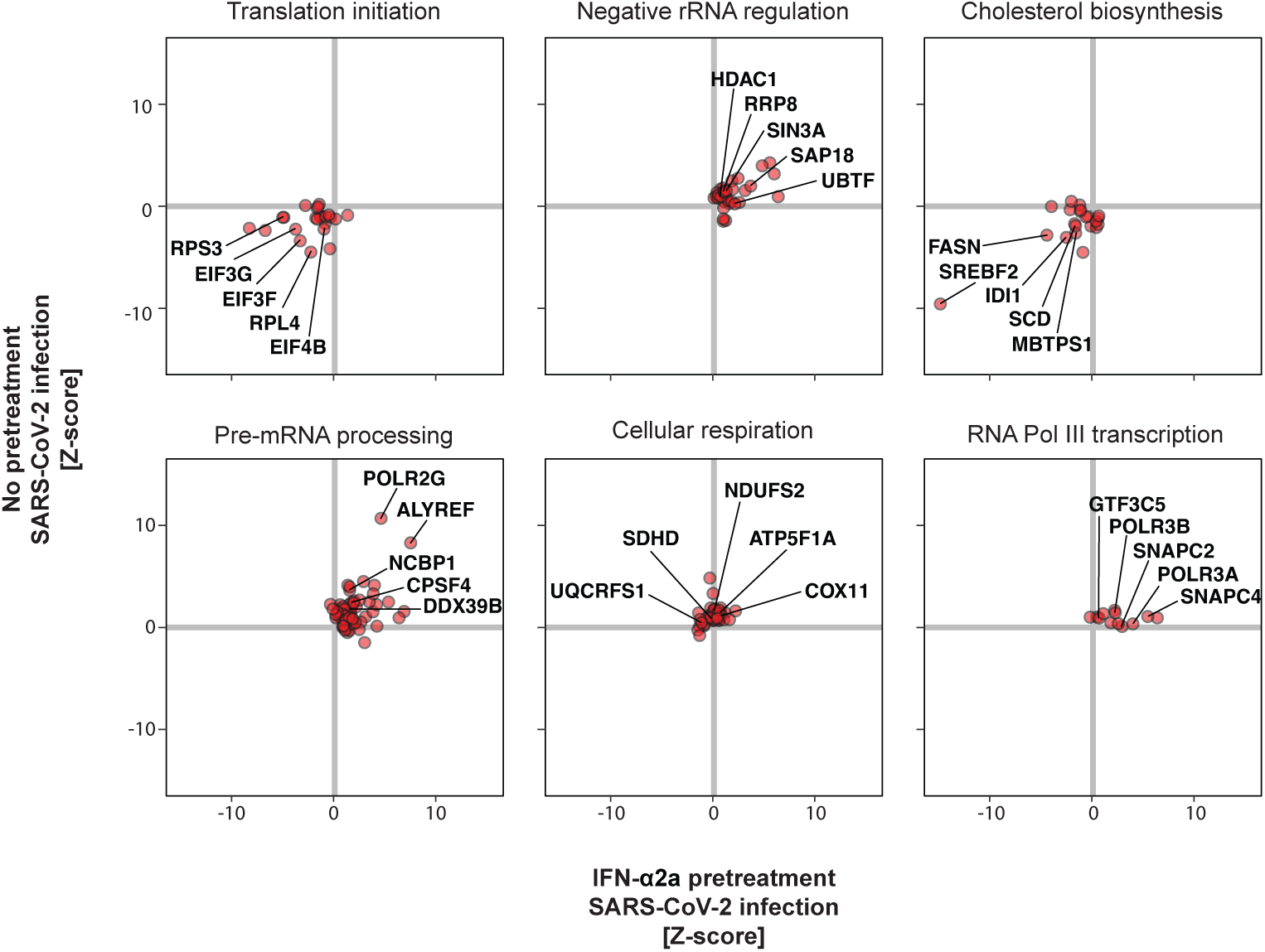
Cellular pathways influencing SARS-CoV-2 infection. GSEA was performed on the screen data (**Supp Table 10**). The individual genes for some of the top pathways are shown here. The full names of the plotted pathways are as follows: translation initiation, “REACTOME EUKARYOTIC TRANSLATION INITIATION”; negative rRNA regulation, “REACTOME NEGATIVE EPIGENETIC REGULATION OF RRNA EXPRESSION”; cholesterol biosynthesis, “REACTOME REGULATION OF CHOLESTEROL BIOSYNTHESIS BY SREBP SREBF”; pre-mRNA processing, “REACTOME PROCESSING OF CAPPED INTRON CONTAINING PRE MRNA”; cellular respiration, “WP ELECTRON TRANSPORT CHAIN OXPHOS SYSTEM IN MITOCHONDRIA”; RNA Pol III transcription, “REACTOME RNA POLYMERASE III TRANSCRIPTION INITIATION FROM TYPE 3 PROMOTER.” Pathways are from the Reactome [188] and Wikipathways [190] databases. The data underlying this Figure can be found in **Supp Table 1**.

Our arrayed CRISPR KO screen results constitute a valuable resource for research on coronavirus infection specifically, as well as on innate immunity in general. These can be used to help characterize human genes influencing SARS-CoV-2 infection and the IFN response.

### The ISG PLSCR1 is associated with COVID-19 outcomes and exhibits antiviral effects in functional SARS-CoV-2 genetic screens

To provide a thorough perspective on human genes that impact SARS-CoV-2 infection and to place our arrayed CRISPR KO screen results within the context of existing research, we have compiled a table that includes findings from a selection of 67 large-scale ‘omic’ studies related to SARS-CoV-2. This compilation encompasses this study and 25 other functional genetic screens for genes that influence SARS-CoV-2 infection [11–13, 15, 18, 19, 28, 31, 64–80], 24 human genetic studies that correlate certain alleles with severe COVID-19 outcomes [5, 6, 8, 25, 26, 81–99], ten publications detailing SARS-CoV-2 protein interactomes [100–109], six focusing on SARS-CoV-2 RNA interactomes [110–115], and one that examines proteins with altered phosphorylation states in SARS-CoV-2-infected cells [116] **(Supp Tables 11-12 for the full, and summary tables, respectively)**. This table highlights the depth of research in publications addressing SARS-CoV-2 infection: genes reported in several independent large-scale studies are more credible candidates for biological relevance (**Supp Fig 3**). As expected, genes associated with the IFN pathway, such as *IFNAR2*, *OAS1*, and *ZC3HAV1/ZAP*, frequently emerged as significant in SARS-CoV-2 studies.

The overlap of gene hits influencing SARS-CoV-2 between our arrayed screen and fifteen published whole-genome pooled screens [19, 31, 64–67, 69–71, 74, 75, 77–80] was higher than expected by chance (**Supp. Fig 4**). This finding indicates that genetic screens conducted with different methods (e.g., CRISPR activation or CRISPR KO, pooled or arrayed format) and in various cellular contexts (e.g., Huh-7.5, A549-ACE2, Calu-3, with or without IFN pretreatment) exhibit both specificity and significant overlap. A pathway analysis of the hits from our arrayed screen, alongside hits from pooled screens, is available in **Supp Table 13**.

We focused on 28 genes identified in both human genetic studies of COVID-19 patients and in functional genetic screens in cell culture, including our own (**Fig 3**). These genes are likely to have significant physiological relevance and to be well-suited for mechanistic studies in cell culture. Among these, the ISG *PLSCR1* stood out, being identified as one of the most potent antiviral genes in our screen (**Fig 1F**). *PLSCR1* variants have been linked to severe COVID-19 in a recent GWAS (listed in **Table 1**)[25, 26]. This was attributed to a role of PLSCR1 in regulating the IFN response in COVID-19 patients [26]. Indeed, a pioneering study showed that PLSCR1 potentiates the transcriptional response to IFN-β treatment in human ovarian carcinoma Hey1B cells [24]. However, PLSCR1 surprisingly appeared as a potent SARS-CoV-2 antiviral even in the absence of IFN in our screens, suggesting a cell intrinsic, IFN-independent function. In other words, baseline levels of PLSCR1 may be sufficient to restrict SARS-CoV-2, and IFN pretreatment could simply enhance this effect by elevating cellular PLSCR1 levels.

**Figure 3.**
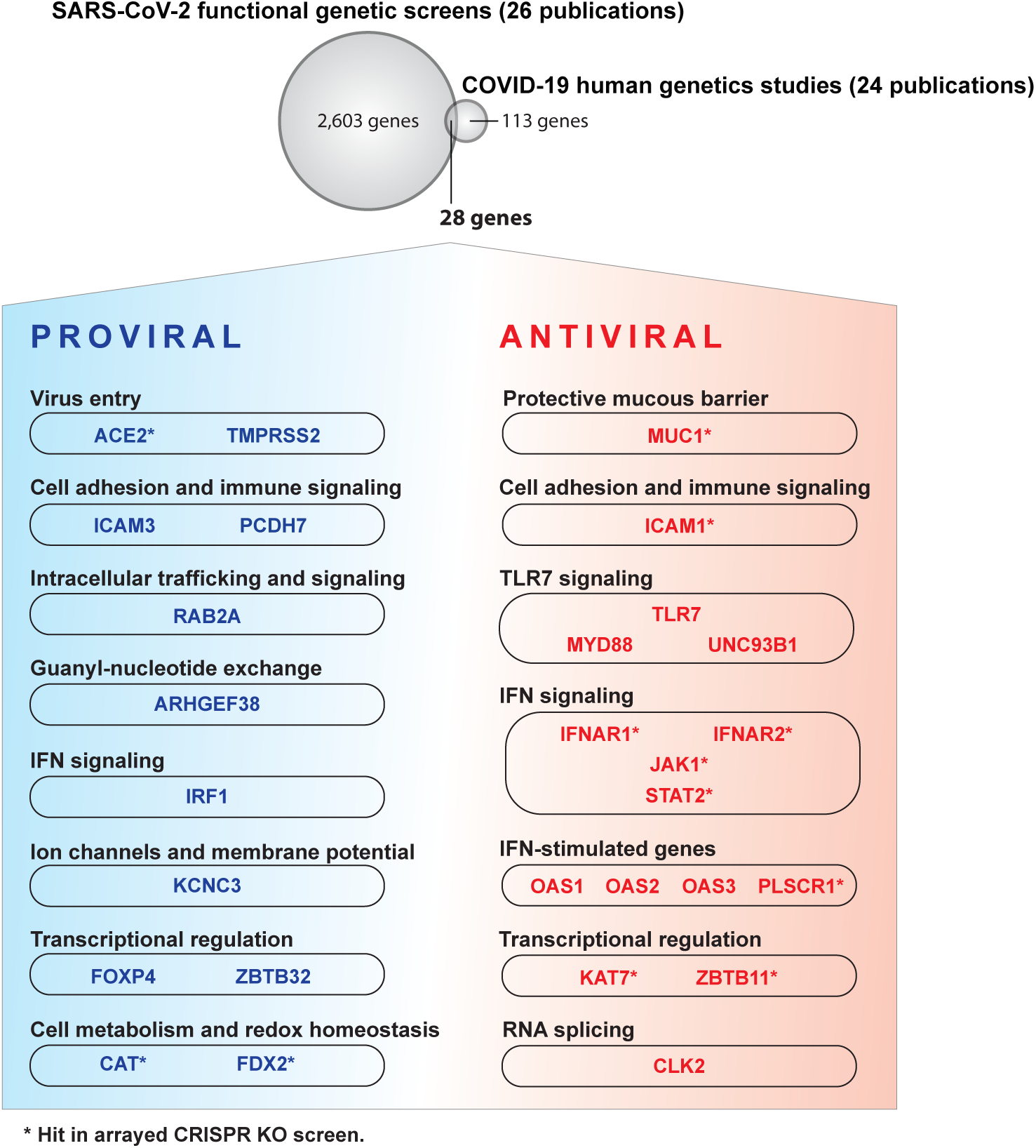
Human genes significant in human genetics studies on COVID-19 patients and in functional genetic screens in cell culture. Overlap between genes considered significant by human genetics studies on COVID-19 patients [5, 6, 8, 25, 26, 81–99] and functional genetic screens for genes influencing SARS-CoV-2 infection in cell culture [11–13, 15, 18, 19, 28, 31, 64–80]. The full list of genes is available in **Supp Table 11**, and a summary is in **Supp Table 12**. Please note that not all genes represented here will influence COVID-19 outcomes. We apologize to the many colleagues whose work was not cited and discussed.

**Table 1.**
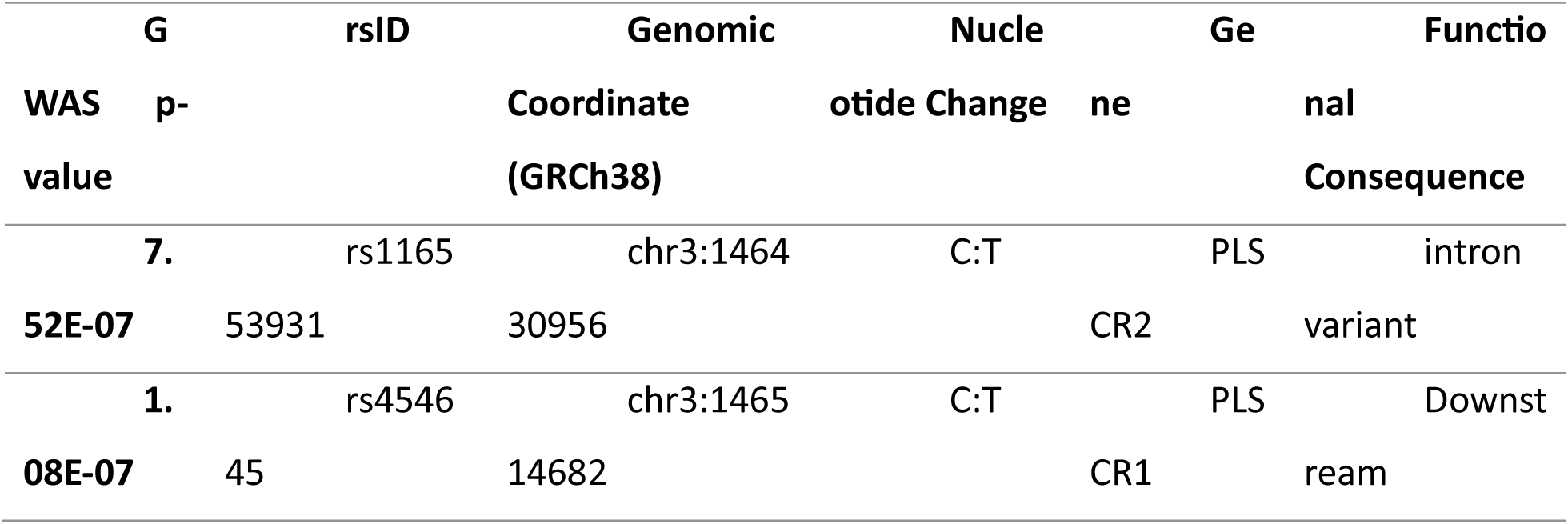
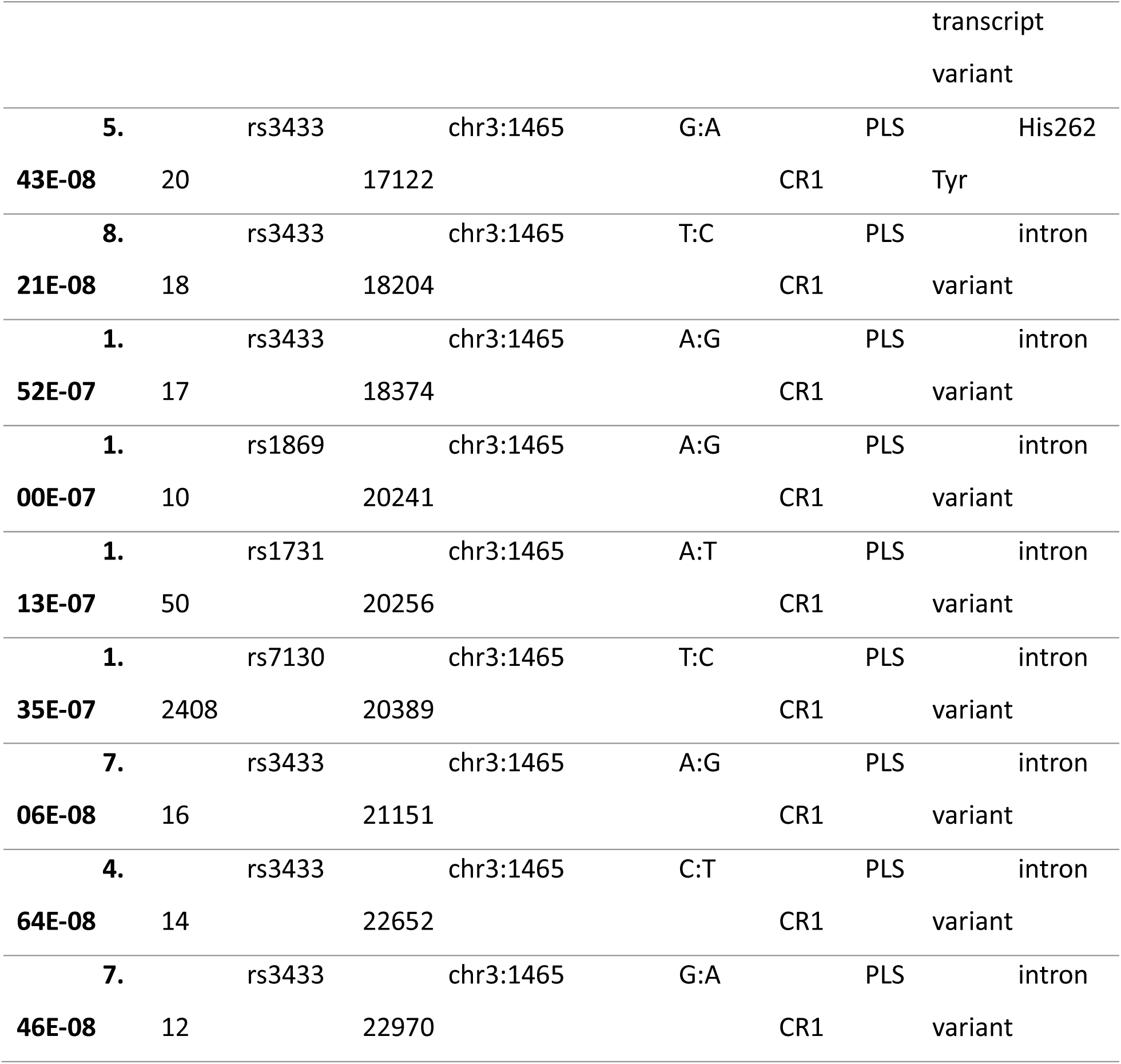
PLSCR1 variants associated with severe COVID-19 in a GWAS [25, 26].

### Intrinsic PLSCR1 restricts SARS-CoV-2 independently of the IFN pathway

To better characterize the function of PLSCR1 during SARS-CoV-2 infection, we generated and validated by western blot (WB) PLSCR1 KO bulk Huh-7.5 and A549-ACE2 lines (**Supp Fig 5A**). As observed in the arrayed screen (**Supp Fig 2B**), PLSCR1 KO cells were viable (**Supp Fig 5B**). PLSCR1 depletion increased susceptibility to SARS-CoV-2 independently of IFN pretreatment (**Fig 4A**). Cell treatment with a JAK-STAT inhibitor, which effectively abrogated IFN signaling, confirmed that intrinsic PLSCR1 limits SARS-CoV-2 infection independently of the IFN signaling pathway (**Fig 4A**). SARS-CoV-2 susceptibility of PLSCR1 KO cells was reversed by the ectopic expression of PLSCR1 (**Fig 4B**, **Supp Fig 5C**). Interestingly, while PLSCR1 tagged with an N-terminal FLAG tag could rescue, PLSCR1 tagged with a C-terminal FLAG tag could not. The C-terminus of the protein is extracellular, and previous research suggests that this region is important for the protein’s scramblase activity and Ca^2+^ binding [117]. It is possible that the addition of this FLAG-tag impaired Ca^2+^ binding, affected PLSCR1’s localization at the plasma membrane, or otherwise disrupted the structure of this region, thereby abolishing PLSCR1’s antiviral ability. We co-cultured PLSCR1 reconstituted cells and PLSCR1 KO cells in the same well and infected them with SARS-CoV-2. A higher proportion of PLSCR1 KO than PLSCR1 reconstituted cells were positive for SARS-CoV-2 indicating that PLSCR1 acts in a cell autonomous manner (**Fig 4C**). Overall, our data indicate that intrinsic PLSCR1 restricts SARS-CoV-2 in cell culture, even in the absence of IFN. Given that PLSCR1 mRNA is constitutively expressed in SARS-CoV-2 target cells (**Supp Fig 5D**) [118], its intrinsic antiviral function may also be effective *in vivo*.

**Figure 4.**
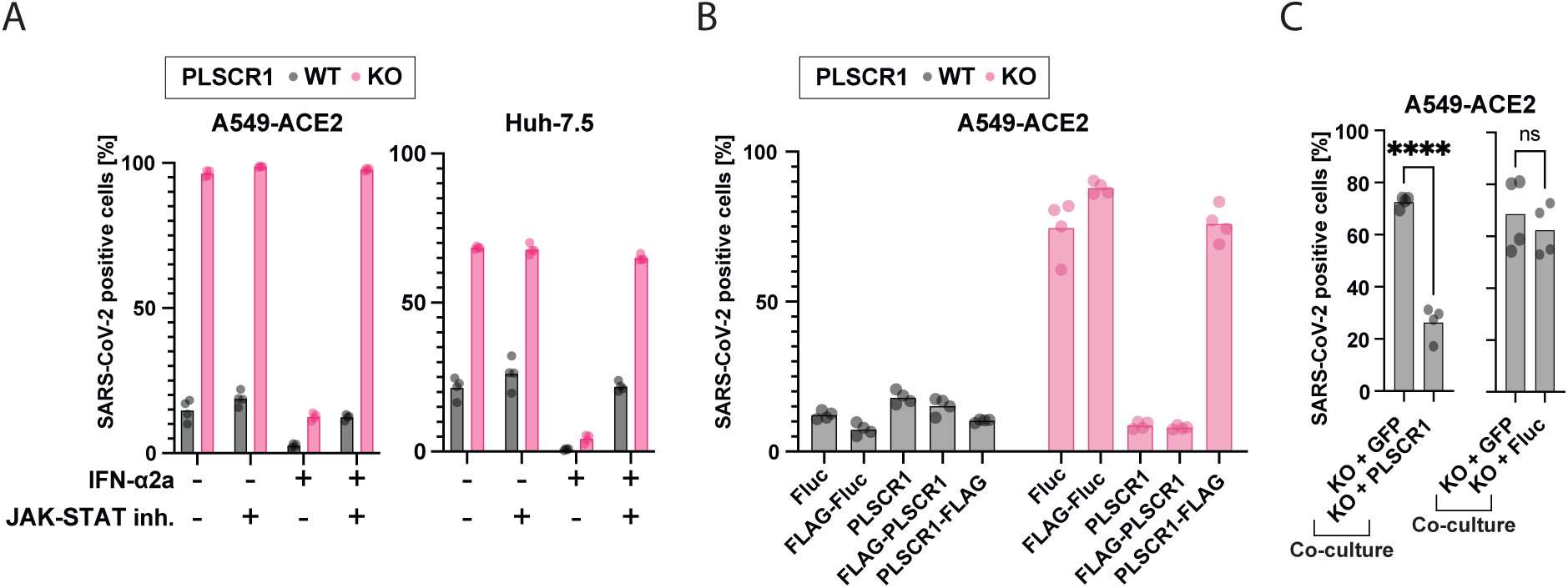
PLSCR1 is a highly effective anti-SARS-CoV-2 effector ISG contributing to intrinsic immunity in the absence of IFN. A. Cells were pretreated with a JAK-STAT inhibitor (InSolution 1 µM) for 2 h, followed by IFN-ɑ2a (10 pM Huh-7.5 or 20 µM A549-ACE2) for 24 h and were infected with SARS-CoV-2 for 24 h followed by IF staining for viral N protein. Huh-7.5 infection using an MOI of 0.5 (titer determined by focus forming assay on Huh-7.5 WT cells). A549-ACE2 infection using an MOI of 0.01 (titer determined by focus forming assay on A549-ACE2 WT cells). The percentage of SARS-CoV-2 positive cells is plotted. n = 4 separate wells infected on the same day. B. Cells were reconstituted with the indicated proteins by stable transduction with lentiviruses, then infected as in **(A)**. n = 4 separate wells infected on the same day. C. Cells were co-cultured as indicated (50:50 mix), then infected as in **(A)** and the % infection of each cell type was determined. n = 4 separate wells infected on the same day; ****, p ≤ 0.0001; two-tailed t-test. The data underlying this Figure can be found in **Supp Table 1**.

### IFN signaling is unaffected by the loss of PLSCR1 in A549-ACE2 and Huh-7.5 cells

PLSCR1 has been shown to potentiate ISG transcription in IFN-treated Hey1B cells [24]. We thus hypothesized PLSCR1 might enhance the type I IFN response in A549-ACE2 and Huh-7.5 cells. We investigated PLSCR1’s role in the IFN response by infecting Huh-7.5 cells with chikungunya virus (CHIKV), which is unaffected by PLSCR1 KO without IFN (**Supp Fig 6A**). PLSCR1 depletion did not functionally affect the antiviral effects of IFN treatment (**Supp Fig 6B**). Furthermore, IFN treatment induced *OAS1* and *IFI6*, two ISGs known to restrict SARS-CoV-2 [11, 13, 18, 95, 96, 98, 99], to a similar extent in both WT and PLSCR1 KO cells, indicating that the IFN signaling pathway was unaffected by PLSCR1 depletion (**Supp Fig 6C-J**). Finally, PLSCR1 depletion did not alter basal ISG transcription in the absence of IFN (**Supp Fig 7, Supp Tables 14 and 15**).

These findings indicate that PLSCR1 limits SARS-CoV-2 infection independently of the IFN signaling pathway in A549-ACE2 and Huh-7.5 cells.

### PLSCR1 restricts SARS-CoV-2 entry

We hypothesized that PLSCR1 directly targets and inhibits a specific step of the SARS-CoV-2 life cycle. PLSCR1 primarily localized at the plasma membrane in Huh-7.5 cells (**Fig 5A**). Furthermore, PLSCR1 depletion led to increased SARS-CoV-2 foci formation (**Fig 5B,C**), and PLSCR1 KO cells did not show increased susceptibility to a SARS-CoV-2 replicon system that bypasses entry (**Fig 5D**) [119]. In contrast, a single-cycle, replication-defective human immunodeficiency virus type-1 (HIV-1) particles pseudo-typed with SARS-CoV-2 spike showed enhanced entry in PLSCR1 depleted cells (**Fig 5E, F**) [120]. This data indicates that PLSCR1 restricts SARS-CoV-2 spike-mediated virion entry. Over-expression of TMPRSS2 lifted the PLSCR1-mediated restriction of authentic SARS-CoV-2 (**Fig 5G-I**) and of SARS-CoV-2 spike pseudo-typed particles (**Fig 5J-M**), indicating that PLSCR1 primarily restricts the endosomal entry route.

**Figure 5.**
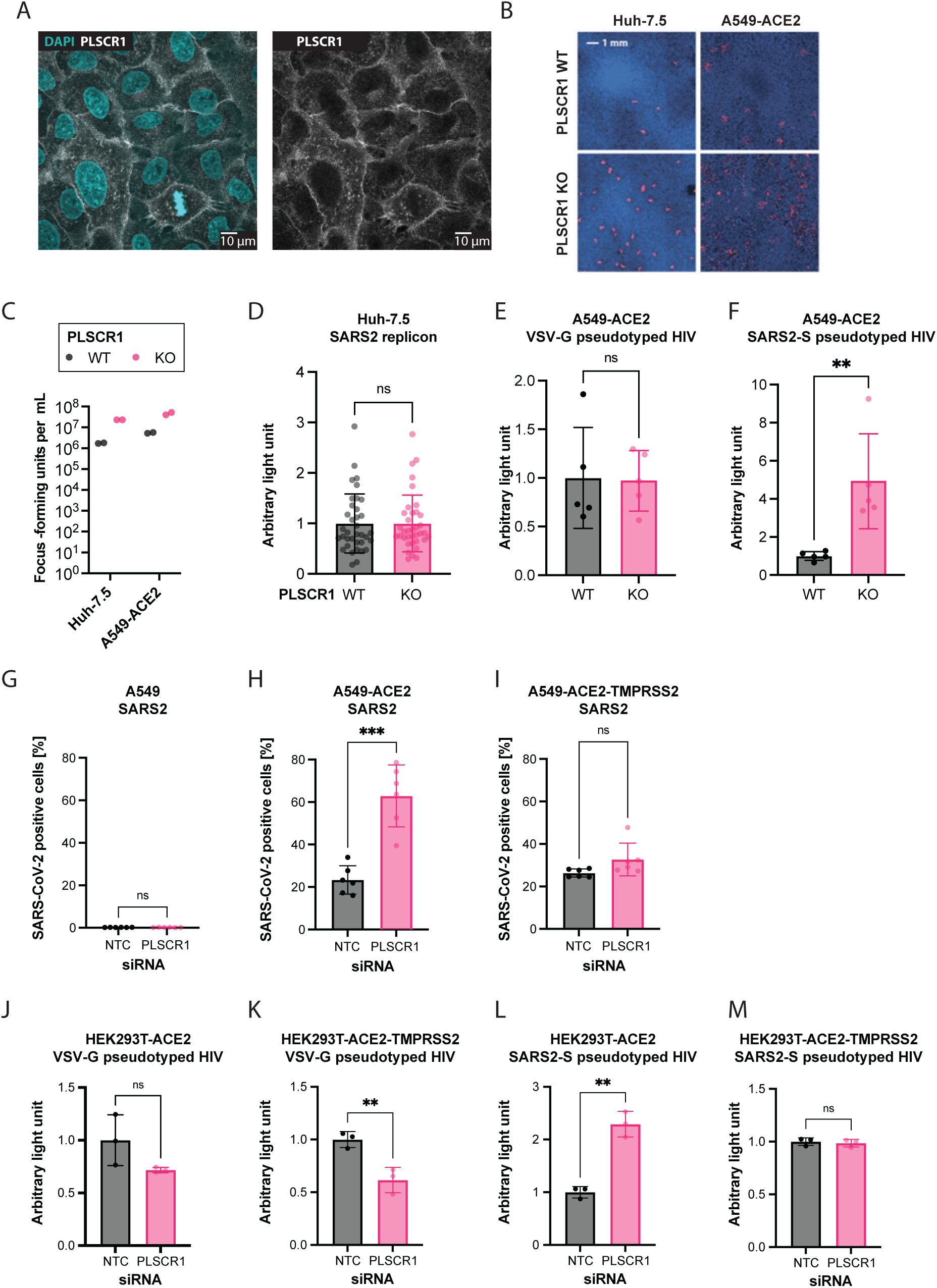
PLSCR1 restricts spike-mediated SARS-CoV-2 entry. A. A549-ACE2 cells were IF stained using an anti-PLSCR1 antibody (white) and Hoechst 33342 nuclear staining (blue) and imaged at 63 X magnification on a confocal microscope. B. Focus forming assays: SARS-CoV-2 N IF (red) and Hoechst 33342 nuclear staining (blue) on similarly infected WT or PLSCR1 KO Huh-7.5 and A549-ACE2 cells after 2 and 3 d, respectively. C. Quantification of **(B)**. D. Huh-7.5 WT and PLSCR1 KO cells electroporated with SARS-CoV-2 replicon which produces a secreted luciferase. Luciferase activity assayed 24 hours after electroporation. n = 36 separate wells from a single electroporation event. Error bars represent sd; ns = non-significant; two-tailed t-test. E-F. Transduction of A549-ACE2 cells with an HIV-based replicon expressing the nanoluciferase pseudotyped with VSV-G or SARS-CoV-2 spike, respectively. n = 5 separate wells transduced on the same day. Nanoluciferase signal measured 2 dpi. Error bars represent sd; ns, non-significant; **, p ≤ 0.01; two-tailed t-test. G-I. A549 cells WT, expressing ACE2, or expressing ACE2-TMPRSS2 as indicated were transfected with PLSCR1 or non-template control (NTC) siRNAs as indicated for 3 d and infected with SARS-CoV-2 for 1 d. SARS-CoV-2 N was stained by IF and the percentage of positive cells was determined by imaging. n = 6 separate wells infected on the same day. Error bars represent sd; ns, non-significant; **, p ≤ 0.01; ***, p ≤ 0.001; two-tailed t-test. J-M. HEK293T cells expressing ACE2 or ACE2-TMPRSS2, as indicated, were transfected with siRNA knockdown of PLSCR1 or non-template control (NTC), as indicated, for 3 d and transduced with an HIV-based replicon expressing the nanoluciferase pseudotyped with VSV-G or SARS-CoV-2 spike, as indicated for 2 d. n = 3 separate wells transduced on the same day. Error bars represent sd; ns, non-significant; **, p ≤ 0.01; two-tailed t-test. The data underlying this Figure can be found in **Supp Table 1**.

Other ISG products have been described to restrict SARS-CoV-2 entry (**Supp Fig 1**) [1], notably: CH25H promotes cholesterol sequestration in lipid droplets, decreasing the pool of accessible cholesterol required for virus-cell membrane fusion [28,29]; LY6E blocks virus-cell membrane fusion via currently undetermined mechanisms [121]; nuclear receptor coactivator 7 (NCOA7) over-acidifies the lysosome, leading to viral antigen degradation by lysosomal proteases [31,32]; and interferon induced transmembrane protein 2 (IFITM2) blocks pH- and cathepsin-dependent SARS-CoV-2 virus-cell membrane fusion in the endosome [122]. The aforementioned ISGs were not hits in our CRISPR KO screen for antiviral genes in IFN-treated cells. This may be due to functional redundancies among them. We found that over-expression of any of the aforementioned ISGs restricted SARS-CoV-2 in PLSCR1 KO cells, indicating that they do not need PLSCR1 for their antiviral activity (**Supp Fig 8A**). PLSCR1 is unlikely to require CH25H, LY6E, or IFITM2 for its function as these are expressed at minimal levels in Huh-7.5 cells without IFN pre-treatment (**Supp Fig 8B,C, Supp Tables 4, 5, 16, and 17**). We cannot exclude an association between PLSCR1 and NCOA7, although the latter was not hit in our screen.

In addition to SARS-CoV-2, we also evaluated the antiviral activity of PLSCR1 against ten viruses that utilize endosomal entry: CHIKV, human parainfluenza virus (hPIV), herpes simplex virus 1 (HSV-1), influenza A virus (IAV), human coronavirus OC43 (hCoV-OC43), human coronavirus NL63 (hCoV-NL63), human coronavirus 229E (hCoV-229E), Sindbis virus (SINV), Venezuelan equine encephalitis virus (VEEV), and vesicular stomatitis virus (VSV). Only SARS-CoV-2 showed a notable susceptibility to PLSCR1’s inhibitory effects (**Supp Fig 9**).

One hypothesis for PLSCR1’s specificity for SARS-CoV-2 is that it alters the surface levels of its receptor, ACE2. However, flow cytometry on live cells did not show a significant effect of PLSCR1 on ACE2 surface levels in A549-ACE2 cells (**Supp Fig 10**). The precise mechanism of action of PLSCR1 remains undetermined.

### Recent variants of SARS-CoV-2 are less restricted by PLSCR1

During the COVID-19 pandemic, SARS-CoV-2 variants evolved from the initial strain, showing increased immune evasion and transmissibility [123–125]. To examine if these variants could circumvent the antiviral action of PLSCR1, we infected WT and PLSCR1 KO Huh-7.5 cells with an early strain isolated in July 2020 (NY-RU-NY1, subsequently referred to as “parental”) and the Beta (B.1.352), Delta (B.1.617.2), Omicron (BA.5), and Omicron (XBB.1.5) variants. PLSCR1 continued to restrict these later variants when examining the percentage of infected cells 24 hours post-infection (**Figs 6A-E**).

**Figure 6.**
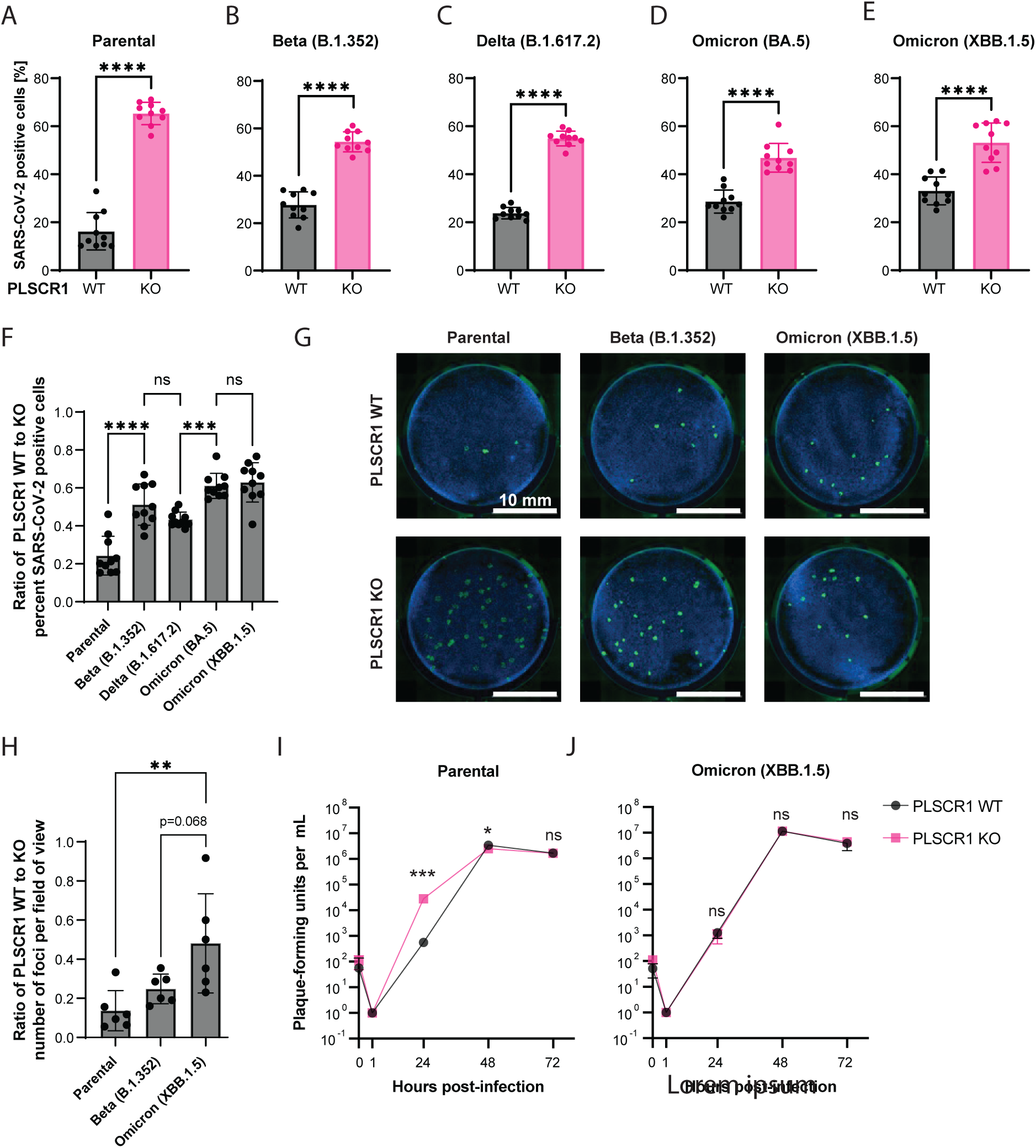
Newer variants of SARS-CoV-2 are less restricted by PLSCR1. A-E. Infection of Huh-7.5 cells with SARS-CoV-2 (parental) or its descendant variants, Beta, Delta, Omicron BA.5, and Omicron XBB.1.5 for 24 hours. SARS-CoV-2 N was stained by IF and the percentage of positive cells determined by imaging. n = 10 separate wells infected on the same day. Error bars represent sd; ****, p < 0.0001; two-tailed t-test. F. Ratio of WT/KO percent infection from A-E. Error bars represent sd. ns, non-significant; ***, p ≤ 0.001; ****, p ≤ 0.0001; one-way ANOVA. G. Focus forming assay on Huh-7.5 WT and PLSCR1 KO cells infected with approximately 50 FFU of SARS-CoV-2 variants as indicated. FFUs were determined on PLSCR1 KO Huh-7.5 cells. Representative images. H. Foci from (**G**) were counted, and then a ratio of WT-to-KO plotted for each SARS-CoV-2 variant. n = 6 separate wells infected on the same day. Error bars represent sd. **, p ≤ 0.01; one-way ANOVA. I. Virus production over an infectious time course (growth curve) for the parental SARS-CoV-2 strain. n = 2 to 3 separate wells infected on the same day. Error bars represent sd. ns, non-significant, *, p ≤ 0.05, ***, p ≤ 0.001, two-tailed t-test on the log10-transformed values. J. As in (**I**) with the Omicron (XBB.1.5) strain. The data underlying this Figure can be found in **Supp Table 1**.

To determine if the magnitude of PLSCR1 restriction was the same for the parental SARS-CoV-2 strain and recent variants, we plotted the percentage of infection data from **Figs 6A-E** as a ratio of PLSCR1 WT to KO (**Fig 6F**). Recent SARS-CoV-2 variants showed reduced differences in infection rates between PLSCR1 WT and KO cells than the parental SARS-CoV-2 strain. The diminished difference in sensitivity between PLSCR1 WT and KO cells was most pronounced with Omicron BA.5 and its descendant, XBB.1.5 (**Fig 6F**).

To examine this further, we infected PLSCR1 WT and KO Huh-7.5 cells with approximately 50 focus-forming units (FFU) per well for different SARS-CoV-2 strains (**Figs 6G and 6H**). In line with **Fig 6F**, the data indicate that the difference in virus susceptibility between PLSCR1 WT and KO cells is lower for more recent variants such as Beta (B.1.352) and especially Omicron (XBB.1.5) compared with the parental strain (**Fig 6H**).

Finally, we quantified virus production over an infectious time course for the parental SARS-CoV-2 strain versus Omicron (XBB.1.5) in Huh-7.5 PLSCR1 WT and KO cells (**Figs 6I and J**). As expected, parental SARS-CoV-2 replicated with faster kinetics upon PLSCR1 depletion. In contrast, Omicron (XBB.1.5) replication was not affected by PLSCR1 depletion. Our data suggest that PLSCR1 restricts newer SARS-CoV-2 variants less efficiently in Huh-7.5 cells. This could be due to adaptation of the recent variants to directly antagonize PLSCR1 and/or utilize an alternative entry route in Huh-7.5 cells that is TMPRSS2-independent and invulnerable to PLSCR1.

### Association between PLSCR1 variants and severe COVID-19

PLSCR1 encodes a 318 amino acid protein containing a palmitoylation motif and a transmembrane domain which regulate its plasma membrane localization, and a nuclear localization signal (NLS) and transcriptional activation domain thought to be important for its nuclear functions (**Fig 7A**)[22, 126–131].

**Figure 7.**
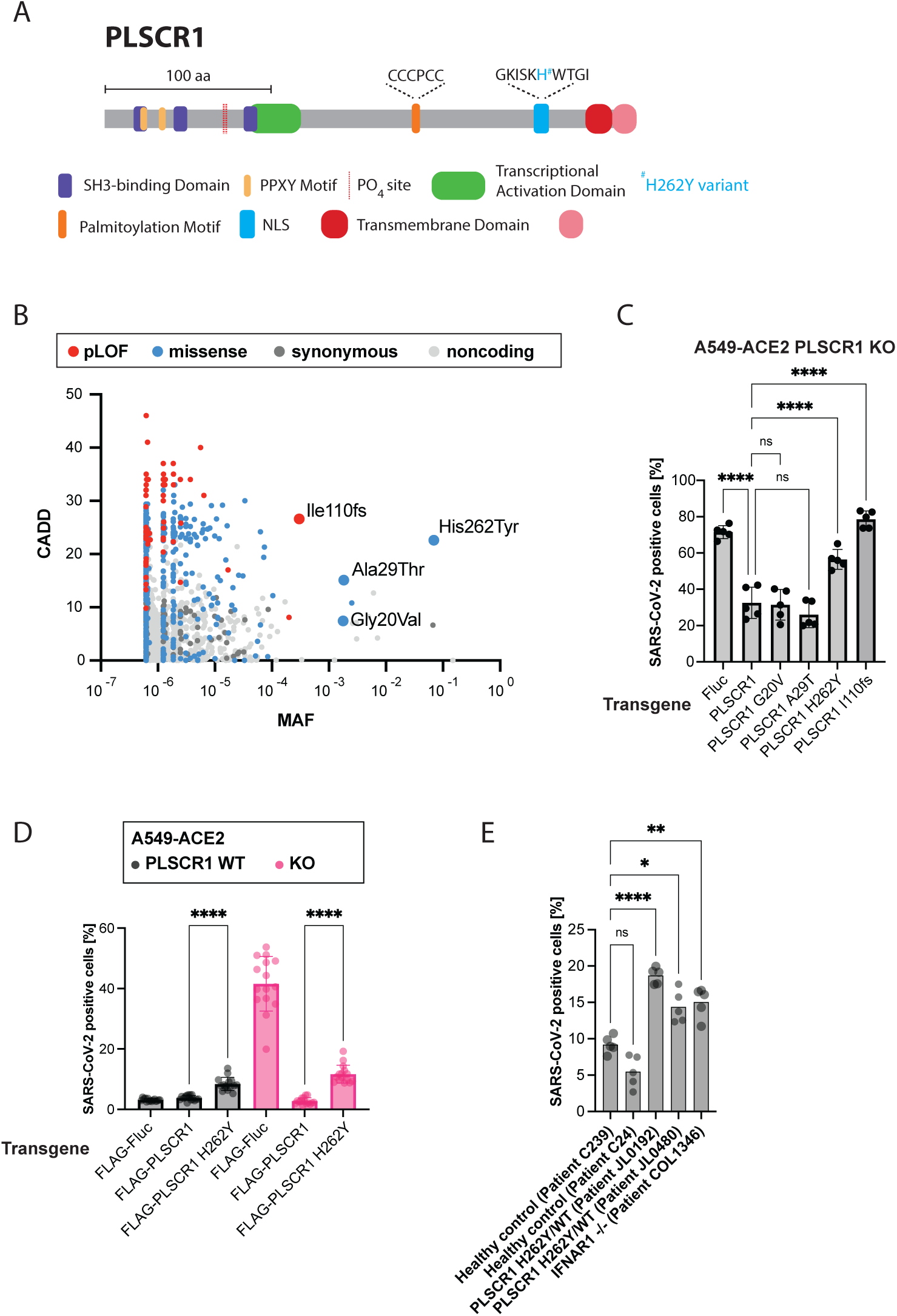
PLSCR1 p.His262Tyr, which associates with severe COVID-19, leads to higher SARS-CoV-2 infection in cell culture. A. Protein diagram of PLSCR1. Domain coordinates from UniProt [197]. B. CADD MAF plot showing the common variants of the PLSCR1 gene reported in gnomAD [132, 133]. The plot displays the Combined Annotation Dependent Depletion (CADD) scores, indicating the predicted deleteriousness of each variant, against the Minor Allele Frequency (MAF). Fs, frameshift. C. A549-ACE2 PLSCR1 KO cells were transduced to stably and ectopically express the indicated PLSCR1 variants. The cells were then infected for 24 h with SARS-CoV-2. SARS-CoV-2 N was stained by IF and the percentage of positive cells determined by imaging. n = 4 separate wells infected on the same day. Error bars represent sd; **, p ≤ 0.01; ****, p < 0.0001; one-way ANOVA. Fs, frameshift. D. A549-ACE2 cells, WT and PLSCR1 KO, stably expressing N-terminal FLAG-tagged Firefly Luciferase (Fluc), N-terminal FLAG-tagged PLSCR1, or N-terminal FLAG-tagged PLSCR1 H262Y mutant and infected with SARS-CoV-2 for 24 hours. SARS-CoV-2 N was stained by IF and the percentage of positive cells determined by imaging. n = 15 separate wells infected on the same day. Error bars represent sd; ****, p ≤ 0.0001; two-tailed t-test. E. SV40-Fibroblast-ACE2 cells, genotype as indicated, infected for 24 hours with SARS-CoV-2. n = 8 separate wells infected on the same day. ns, non-significant; *, p ≤ 0.05; **, p ≤ 0.01; ****, p ≤ 0.0001; one-way ANOVA. The data underlying this Figure can be found in **Supp Table 1**.

A recent GWAS has identified an association between PLSCR1 variants and severe COVID-19 outcomes, reporting an odds ratio of approximately 1.2 and a p-value of approximately 10^-8^ (**Table 1**)[25, 26]. In other words, the GWAS suggests that PLSCR1 has a small, but significant effect on severe COVID-19 risks.

GWAS typically identify variants associated with increased odds of a disease, but these variants are not necessarily causative. Among the PLSCR1 variants identified in the COVID-19 GWAS cited above, only p.His262Tyr (rs343320) results in a protein-coding change, located in the NLS (**Fig 7A**, **Table 1**). This raises the hypothesis that p.His262Tyr might alter PLSCR1’s antiviral function. However, we cannot dismiss the possibility that (i) some non-coding variants identified in the GWAS could influence the regulation of PLSCR1 mRNA, potentially leading to functional outcomes, and (ii) the GWAS might have missed other nonsynonymous variants besides p.His262Tyr that could potentially impact PLSCR1 function.

To assess the relative importance of the p.His262Tyr variant in humans, we examined the full collection of PLSCR1 variants reported in gnomAD, a human population reference database (**Fig. 7B**) [132, 133]. The p.His262Tyr variant is the most common nonsynonymous variant in this gene, with a minor allele frequency (MAF) of 0.07, MAFmax of 0.1, and 4,348 homozygous individuals in gnomAD. Another variant, resulting in a frameshift mutation, p.Ile110fs (rs749938276), is the most common predicted loss-of-function (pLOF) of PLSCR1 in gnomAD (MAF of 0.0003, MAFmax of 0.01, 1 homozygous individual), indicating that PLSCR1 is not essential.

We aimed to investigate the functional effects of the p.His262Tyr and p.Ile110fs variants in cell culture, along with two other relatively common nonsynonymous variants, p.Gly20Val (rs79551579, MAF of 0.002, MAFmax of 0.04) and p.Ala29Thr (rs41267859, MAF of 0.01, MAFmax of 0.04). For this, we stably and ectopically expressed the PLSCR1 variants of interest in PLSCR1 KO A549-ACE2 cells using a lentiviral vector [134]. The variant p.Ile110fs did not rescue PLSCR1 KO, confirming it as a loss-of-function mutation (**Fig 7C**). The p.His262Tyr variant, expressed at similar levels to WT PLSCR1 in this system (**Supp Fig. 11A**), behaved as a hypomorphic allele, providing only partial rescue of PLSCR1 antiviral function (**Figs. 7C and D**). Additionally, introducing PLSCR1 p.His262Tyr into PLSCR1 WT A549-ACE2 cells increased their susceptibility to SARS-CoV-2 infection (**Fig 7D**), suggesting a dominant effect. However, this effect might be attributed to the overexpression of PLSCR1 p.His262Tyr from the transgene, compared to the natural expression levels of PLSCR1 WT from the endogenous locus. To counter this, we examined patient-derived SV40-immortalized fibroblasts expressing ACE2 that were heterozygous for p.His262Tyr. These cells were already present in our collection, originating from a female tuberculosis patient from Turkey (JL0192) and a female herpes simplex encephalitis patient from France (JL0480). The origin of these cells is serendipitous; our goal was not to link these diseases to PLSCR1 but rather to infect cells carrying the His262Tyr variant with SARS-CoV-2. Cells heterozygous for p.His262Tyr were hyper-susceptible to SARS-CoV-2 infection compared to PLSCR1 WT control SV40-fibroblasts (**Fig 7E**), further suggesting that p.His262Tyr is dominant. We cannot formally rule out that the examined SV40-fibroblasts may carry other mutations influencing SARS-CoV-2 infection.

Since the p.His262Tyr mutation is located in the NLS region, we hypothesized that it could impair PLSCR1’s nuclear localization. However, PLSCR1 WT was primarily localized in the cytoplasm in untreated A549-ACE2 cells (**Fig 5A**), in IFN-treated cells (**Supp Fig 11B**), and in SARS-CoV-2 infected cells (**Supp Fig 11C**). Similarly, PLSCR1 His262Tyr was also enriched in the cytoplasm (**Supp Fig 12**). This suggests that the palmitoylation motif, known to be dominant over the NLS [129], dictates the cytoplasmic localization of PLSCR1. Therefore, we propose that the p.His262Tyr mutation affects PLSCR1 through a mechanism other than altering its nuclear localization.

Our data collectively highlight PLSCR1’s function in restricting SARS-CoV-2 entry in cell culture, thereby clarifying the association between PLSCR1 variants and severe COVID-19 outcomes [25, 26]. Future human genetic studies are crucial for determining if certain PLSCR1 variants in the population cause increased risks of severe COVID-19. Our results highlight the variant p.His262Tyr as a potential causative candidate, as it caused increased SARS-CoV-2 infection in cell culture.

## Discussion

Here, we conducted an unbiased arrayed CRISPR KO screen on Huh-7.5 cells infected with SARS-CoV-2. The screen revealed novel aspects of SARS-CoV-2 and IFN biology while also confirming previously known facets. Pathways related to mRNA transcription and maturation were identified as antiviral. This observation may stem from the conflict between the host cell and SARS-CoV-2, where the host attempts to export mRNAs from the nucleus to facilitate antiviral responses while the virus replicates in the cytoplasm, impeding nuclear export [135–138]. RNA Pol III transcription was specifically essential for the IFN-mediated antiviral response, through mechanisms that are yet to be determined. Interestingly, inborn errors in POLR3A and POLR3C have been previously described in patients with severe varicella zoster virus infections [139]. Cellular respiration was identified as a key IFN-independent antiviral pathway. Furthermore, mitophagy was identified as proviral. This may indicate the infected cell’s increased demand for energy and ATP to combat the virus. Alternatively, cellular respiration may have other, yet-to-be-identified, IFN-independent antiviral roles. Conversely, translation and cholesterol homeostasis emerged as the foremost proviral pathways. These findings underscore the complex, dualistic nature of the interactions between SARS-CoV-2 and host cells.

Our screen notably identified the ISG zinc-finger antiviral protein (*ZC3HAV1*/*ZAP*) as a proviral factor in IFN-treated cells. Initially, ZC3HAV1/ZAP gained attention as an antiviral factor that targets the SARS-CoV-2 RNA genome [115] and prevents programmed ribosomal frameshifting [140]. Yet, a recent study demonstrated that ZC3HAV1/ZAP also promotes the formation of SARS-CoV-2 non-structural proteins 3 and 4-induced double-membrane vesicles, essential for virus replication [72]. SARS-CoV-2 may have adapted to exploit certain ISG products, such as ZC3HAV1/ZAP, within the cellular environment it encounters. It is still unclear if the seemingly contradictory roles of ZC3HAV1/ZAP – both proviral and antiviral – are caused by distinct isoforms.

Many other ISG products influenced SARS-CoV-2 infection, PLSCR1 being the most potent restriction factor. PLSCR1 did not influence ISG induction as previously reported [24], but rather inhibited spike-mediated SARS-CoV-2 entry through the endocytic route. Our results corroborate a recent study from Xu et al [19] and provide an explanation for the enrichment for PLSCR1 SNPs observed in a GWAS on severe COVID-19 [25, 26].

The molecular mechanisms of PLSCR1-mediated restriction of SARS-CoV-2 entry remain to be elucidated. PLSCR1 could be altering the lipid composition at the contact site between the virus and endosomal membranes, akin to the ISG interferon induced transmembrane protein 3 (IFITM3) for influenza A virus [141–144]. PLSCR1 was first identified as a Ca^2+^-dependent phospholipid scramblase [23], but it is unclear whether PLSCR1 depletion affects the bidirectional movement of phospholipids *in vivo* [145–147]. A C-terminal FLAG-tag abolished the antiviral ability of reconstituted PLSCR1, possibly by interfering with the function of the Ca^2+^ binding domain. In contrast, inhibiting PLSCR1’s phospholipid scramblase activity did not alleviate SARS-CoV-2 restriction [19].

PLSCR1 is one of a few ISGs known to restrict SARS-CoV-2 entry (**Supp Fig 1A**)[1, 148], with possibly more yet to be discovered. Future research should investigate the associations and combinatorial effects between antiviral ISGs, which could lead to key mechanistic insights into their mode of action.

Intriguingly, PLSCR1 specifically restricted SARS-CoV-2 in Huh-7.5 and A549-ACE2 cells but it did not show similar inhibitory effects on other viruses that enter cells via endocytosis. Future studies will investigate the mechanisms behind this specificity. PLSCR1 has been described to inhibit a range of viruses in various cell lines, such as encephalomyocarditis virus, vesicular stomatitis virus, Epstein-Barr virus, hepatitis B virus, hepatitis C virus, human cytomegalovirus, human immunodeficiency virus 1, human T-cell lymphotropic virus type 1, and influenza A virus [24, 149–155]. It has been proposed that PLSCR1 directly binds viral proteins and impairs their functions to restrict the non-coronaviruses cited above, reviewed in [156]. However, it seems unlikely that PLSCR1 has evolved to interact directly with such a diverse set of viral proteins. An alternative explanation is that diverse viral proteins convergently evolved to bind PLSCR1 as a mechanism of immune evasion. For example, by altering PLSCR1’s subcellular localization. Meanwhile, overexpressing PLSCR1 in cell culture could act *like a sponge*, absorbing these viral proteins and thereby hindering viral function. We searched for PLSCR1 interactions in ten SARS-CoV-2 proteins interactome studies, relying on ectopic expression of individual viral proteins [100–102, 104–109, 157], and no interaction was reported in two or more independent studies. Two interactions were reported in a single study: (i) PLSCR1-ORF7b [102], and (ii) PLSCR1-ORF8 [105], both by proximity biotinylation, which was less stringent compared to affinity purification-mass spectrometry (AP-MS) and yeast two-hybrid (Y2H) techniques (**Supp Fig 13**). To date, there is no strong evidence of a direct interaction between a SARS-CoV-2 protein and PLSCR1, although we cannot rule out that such interactions may occur or even appear in the future as SARS-CoV-2 evolves.

Previous research suggests that Omicron has developed increased resistance to IFN [158, 159], a trait associated with its highly-mutated spike protein [160]. Here, we show that recent SARS-CoV-2 variants, including Omicron BA.5 and XBB.1.5, exhibit reduced sensitivity to PLSCR1-mediated restriction compared to the New York 2020 strain, which served as a reference in our study (**Fig 6**). Evasion from PLSCR1 may thus have contributed to Omicron’s increased resistance to IFN.

The original SARS-CoV-2 strain can enter cells through both TMPRSS2-dependent fusion near the cell surface and clathrin-mediated endocytosis, with a preference for the former *in vivo* [38]. ISGs that primarily restrict SARS-CoV-2 endocytosis, such as NCOA7, which perturbs lysosome acidification [161, 162], and PLSCR1, may play a role in constraining the original strain to TMPRSS2-dependent entry. Interestingly, the more recent Omicron variant shows both increased evasion from PLSCR1 restriction (**Fig 6**) and an acquired preference for TMPRSS2-independent entry. This occurs either via endocytosis or TMPRSS2-independent fusion near the cell surface, utilizing metalloproteases to cleave the spike protein [160, 163, 164]. We postulate that Omicron evolved its entry pathway to evade PLSCR1-mediated restriction while shifting its cell tropism to the upper airways and away from TMPRSS2-expressing cells [163, 165–169].

Future research should investigate: (i) mechanism(s) by which Omicron evades PLSCR1 restriction and whether this evasion is primarily due to mutations in the spike protein, (ii) the association between PLSCR1-mediated restriction and the distinct entry mechanisms utilized by SARS-CoV-2 variants, (iii) how various intrinsic factors that restrict virus entry, such as PLSCR1, have influenced SARS-CoV-2 entry routes, evolution, and cell tropism. Understanding these aspects of PLSCR1 restriction could provide mechanistic insight into broad strategies of PLSCR1 evasion employed by both newer SARS-CoV-2 variants as well as viruses from other families that were unaffected by PLSCR1 KO. Subversion of PLSCR1 restriction may serve as a useful immune evasion strategy that could be shared by diverse viruses with longer evolutionary selection than SARS-CoV-2.

Several PLSCR1 variants were enriched in a GWAS on severe COVID-19, with a relatively low odds ratio of approximately 1.2 [25, 26]. This modest odds ratio likely reflects the complex redundancies within antiviral defenses, from innate immunity featuring multiple effector ISGs that restrict SARS-CoV-2 [10–15, 18, 19], to adaptive immunity [170]. For example, we show that the ISGs CH25H [28,29], IFITM2 [122], LY6E [121], and NCOA7 [31,32] still function to restrict SARS-CoV-2 in PLSCR1-depleted cells (**Supp Fig 8**). However, for these very reasons, the identification of PLSCR1 variants in the GWAS remains noteworthy. Of these enriched variants, only PLSCR1 p.His262Tyr (rs343320) resulted in a protein-coding change. Our findings indicate that p.His262Tyr exhibits a hypomorphic and dominant effect in cell culture, leading to increased SARS-CoV-2 infection. Future research should aim to ascertain whether p.His262Tyr, or potentially other PLSCR1 variants, are directly responsible for elevated risks of severe COVID-19 in patients.

Our findings show that baseline levels of PLSCR1 are effective in limiting SARS-CoV-2 infection. This is in line with other studies where ISGs like *DAXX* and *LY6E* were shown to inhibit SARS-CoV-2 independently of IFN [18, 121, 171]. mRNA-seq analyses of Huh-7.5 cells and primary human hepatocytes, as well as data from the GTEx consortium on various human tissues [49, 172], revealed that many ISGs are constitutively expressed, even without IFN stimulation (**Supp Fig 14**). This supports the idea that the IFN-induced antiviral state results more from enhanced expression of antiviral genes rather than a binary ON/OFF switch. In future studies, it will be interesting to explore whether intrinsically expressed ISGs also carry out cellular functions beyond pathogen defense.

## Materials and Methods

### Plasmids, oligos, and primers

The plasmids, gene fragments, and primers used in this study are listed in **Supp Tables 18, 19, and 20**, respectively.

### Cell Lines

Huh-7.5 (human hepatocellular carcinoma; *H. sapiens*; sex: male) [173], Huh-7.5-Cas9 [31], A549-ACE2 (human lung carcinoma; *H. sapiens*; sex: male; generously provided by the laboratory of Brad R. Rosenberg) [11], A549-ACE2-TMPRSS2 [31], Lenti-X 293T (*H. sapiens*; sex: female: Takara, cat. #632180), Caco2 (*H. sapiens*; sex: male; ATCC, cat. HTB-37), Vero E6 (*Chlorocebus sabaeus*; sex: female; kidney epithelial cells, ATCC cat. #CRL-1586), BHK-21 (*Mesocricetus auratus*; sex: unspecified; kidney fibroblasts) cells, and SV40-Fibroblasts were cultured in Dulbecco’s Modified Eagle Medium (DMEM, Fisher Scientific, cat. #11995065) supplemented with 0.1 mM nonessential amino acids (NEAA, Fisher Scientific, cat. #11140076) and 10% fetal bovine serum (FBS, HyClone Laboratories, Lot. #KTH31760) at 37°C and 5% CO2. All cell lines tested negative for mycoplasma.

### Virus stocks

**CHIKV-181/25-mKate2:** the infectious clone was a kind gift from Mark Heise (University of North Carolina, USA) [not published yet]. 20 µg of infectious clone DNA was linearized with NotI-HF at 37°C overnight. Complete digestion was confirmed by running a sample of the digested DNA on a 1% agarose gel. After confirmation, linearized DNA was cleaned via phenol-chloroform extraction, and then ethanol precipitated. The precipitated DNA was resuspended in 20 µL of RNase-free H_2_O and in vitro transcribed with an SP6 mMessage mMachine In Vitro Transcription Kit (ThermoFisher, cat. AM1340). The generated RNA was electroporated into 1.2 × 10^7^ BHK-21 cells, and the produced virus was harvested once approximately 80% of the electroporated cells lifted or showed signs of cytopathic effects and 100% of the cells were positive for mKate2 signal. The titer of the virus was 8.5 × 10^6^ focus-forming units (FFU) on Huh-7.5 cells.

**hCoV-NL63:** was generously provided by Volker Thiel (University of Bern) and amplified at 33°C in Huh-7.5 cells as in [31].

**hCoV-OC43**: was obtained from ZeptoMetrix (cat. #0810024CF) and amplified at 33°C in Huh-7.5 cells as in [31].

**hPIV3-GFP** [174]: stock (based on strain JS) grown in VeroE6 cells as in [175].

**HSV-1-GFP**: stock made by passage on VeroE6 cells. 2 × 10^7^ cells seeded in a T175 flask were infected at an MOI of 0.01 PFU/ml of HSV-1-GFP virus engineered and provided by Ian Mohr [176]. After a one-hour incubation at 37°C, the inoculum was removed, and 20 ml of DMEM supplemented to contain 10% FBS and NEAA was added. Cells were incubated at 37°C for 24 h or until CPE was evident. Cell supernatant containing progeny virus was harvested and titrated on Vero E6 cells (2.4% avicel, fix 2 dpi) at 2.4 × 10^8^ PFU/ml.

**IAV WSN (H1N1):** was generated in MDCK cells. Cells were inoculated at MOI 0.01 in DMEM supplemented with NEAA, 0.2% BSA, 0.1% FCS, 50 mM Hepes, and 1 µg/ml TPCK-trypsin. Virus-containing culture supernatant was harvested at 52 h post-infection and cleared by centrifugation.

**SARS-CoV-2:** unless otherwise stated, the isolate SARS-CoV-2/human/USA/NY-RU-NY1/2020 was used in this study [177]. The virus was sourced from the saliva of a deidentified patient in New York City, collected on July 28, 2020. Its sequence is publicly accessible (GenBank OM345241). The virus isolate was amplified in Caco-2 cells. The passage 3 stock employed had a titer of 3.4 × 10^6^ PFU/ml, as measured on Vero E6 cells using a 1% methylcellulose overlay, according to previously described methods [178]. The Beta (B.1.351), Delta (B.1.617.2), Omicron BA.5, and Omicron XBB.1.5 variants were obtained from BEI resources (cat. # NR-54008, NR-55611, NR-58616, and NR-59104, respectively), amplified in Vero E6 cells engineered to stably express TMPRSS2, and titer was determined as described above.

**SINV Toto1101** [179]: expressing an nsP3-mScarletI fusion reporter was generated by cloning the sequence encoding mScarletI in frame into a unique SpeI restriction site in the pToto1101 infectious clone plasmid as previously described [180]. *In vitro* transcribed, capped RNA was generated from the pToto1101-nsP3-mScarletI plasmid (Invitrogen mMessage mMachine SP6 kit, AM1340) and electroporated into BHK-J cells, a derivative of BHK-21 cells (ATCC, CCL-10) as previously described [180]. 24 hours post electroporation, centrifuge clarified supernatants were aliquoted and stored at −80C. BHK-J cells were cultured and virus stocks generated in MEM supplemented with 7.5% FBS.

**VEEV-dsEGFP** [16, 181]: the infectious clone plasmid was linearized (MluI) and transcribed *in vitro* using an mMessage mMachine SP6 transcription kit (Ambion). BHK-21 cells were electroporated with viral RNA, and supernatant containing progeny virus was harvested after incubation at 37°C for 30 h or until CPE was evident. Virus was titrated by plaque assay on BHK-21 cells (2.4% avicel, fix 2 dpi). BHK-21: 1.45 × 10^9^ FPU/ml.

**VSV-GFP** [182]: grown in BHK-21 cells as in [175].

**YFV 17D:** was generated via transfection of Huh-7.5 with in vitro transcribed RNA from pACNR-FLYF-17D plasmid as described in [175].

### mRNA-seq

#### mRNA-seq on SARS-CoV-2-infected cells

**Cell culture and infection:** 75,500 Huh-7.5 cells or 150,000 Calu-3 cells were seeded in each well of a 12-well plate with 1 mL media. Media: DMEM with 5% FBS and 1% NEAA for Huh-7.5 cells or EMEM (ATCC, 30-2003) with 10% FBS for Calu-3 cells. The next day, cells were infected by removing 500 µL of media and adding 500 µL of media with SARS-CoV-2 strain USA-WA1/2020 (BEI Resources, NR-52281) at 5,000 PFU/well (virus titer determined in Huh-7.5 cells). After one day, the wells were washed with PBS and cells were harvested in 1 mL TRIzol (Invitrogen, cat. 15596-018). N = 3 replicates (separate wells) per sample.

**RNA extraction:** 2 ml MaXtract High Density tubes (Qiagen, 129056) were centrifugated at 12,000–16,000 x g for 20-30 second centrifugation. A volume of 750 µL TRIzol-prepared sample was combined with 150 µL chloroform in these tubes and hand-shaken vigorously. Phase separation was accomplished by centrifugation at 1500 x g for 5 min at 4°C. The aqueous phase was then mixed with 400 µL ethanol 95-100% in a separate tube. These preparations were then transferred to Zymo Research RNA clean and concentrator-25 kit columns (Zymo Research, cat. R1018) and subjected to multiple wash and centrifugation steps as recommended by the manufacturer. An in-column DNase I treatment was performed using Qiagen DNase (Qiagen, 79254). Finally, RNA was eluted with 50 µL DNase/RNase-Free water and stored at −80°C.

**Sequencing:** Poly-A enriched libraries were made using the TruSeq stranded mRNA LT kit (Illumina, Cat# 20020594) and sequenced on a NovaSeq SP with PE150 read length.

#### mRNA-seq on IFN-treated cells

**Cell culture and treatment: A549 cells**: 300,000 cells were seeded in each well of a 6-well plate with 3 mL of DMEM supplemented with 10% FBS and 1% NEAA. The following day, the media was aspirated and replaced with 2 mL of DMEM with or without 0.5 nM IFN-α2a (PBL, cat. 11101-2), and the cells were incubated at 37°C. After 24 hours, the cells were harvested in 500 µL TRIzol. N = 3 replicates (separate wells treated on the same day) per sample. **Calu-3 cells**: 200,000 cells were seeded in each well of a 12-well plate with 1 mL media. Two days later, the media was replaced with EMEM (ATCC, cat. 30-2003) with 10% FBS with 0.5 IFN-α2a (PBL, cat. 11101-2) and incubated at 37°C. 24 h later, cells were harvested in 500 µL TRIzol. N = 3 replicates (separate wells treated on the same day) per sample. **Huh-7.5 cells**: 75,500 cells were seeded in each well of a 12-well plate with 1 mL media. Two days later, the media was replaced with 1 mL of DMEM with 5%FBS, 1% NEAA with 0.5 nM IFN-α2a (PBL, cat. 11101-2) and incubated at 37°C. 24 h later, cells were harvested in 500 µL TRIzol. N = 3 replicates (separate wells treated on the same day) per sample.

**RNA extraction** as described above.

**Sequencing:** Poly A-enriched libraries were made using the NEBNext Ultra II RNA Library Prep Kit for Illumina (NEB, cat. E7770) and sequenced on a NovaSeq SP with PE150 read length.

#### mRNA-seq on PLSCR1 KO cells

**Cell culture and CRISPR KO:** 30,000 Huh-7.5 cells were seeded in five wells of a 24-well plate with 480 uL media. The cells were reverse transfected with 120 uL of a transfection mixture composed of 250 nM of pooled anti-PLSCR1 or non-targeting Edit-R crRNAs from Horizon Discovery (cat. CM-003729-01-0002, CM-003729-02-0002, CM-003729-03-0002, and CM-003729-04-0002 or U-007501-01-05, U-007502-01-05, U-007503-01-05, and U-007504-01-05, respectively) which had been resuspended with an equimolar amount of Edit-R tracrRNA (Horizon, cat. U-002005-20) and a 1:200 dilution of Dharmafect 4 (Horizon, cat. T-2004-01). The following day, the media was changed, and the cells were progressively scaled up to a 6-well plate over the next 4 days. When the cells were confluent in the 6-well plate, the media was removed from four of the wells. They were then washed with 1x PBS (cat. 14190-144) and lysed with 1 mL TRIzol (Life Technologies, cat. 15596-018) for 5 minutes at room temperature before transferring to an Eppendorf tube and freezing at −80°C to await RNA extraction. The remaining well was lysed with 300 µL of RIPA buffer (Thermo cat. 89900) supplemented with 1x protease inhibitor (Thermo cat. 87786) and 1x EDTA and prepared for western blot as described below, in the “Western Blots” section.

**RNA extraction** as described above.

**Sequencing:** Poly A-enriched libraries were made using the NEBNext Ultra II RNA Library Prep Kit for Illumina (NEB, cat. E7770) and sequenced on a NovaSeq SP with PE150 read length.

#### mRNA-seq analysis

mRNA-seq reads were first quality-filtered and adapter-trimmed using Trim Galore with parameters -q 20 -e 0.1 --length 20 --paired and Cutadapt. Reads were then mapped to the human genome GRCh38 or to a combined SARS-CoV-2 MN985325.1/human genome GRCh38 using STAR [183] with settings including --runThreadN 8 --outFilterMultimapNmax 1 --twopassMode Basic. Feature counting was performed using the featureCounts function from the Rsubread package [184], with strandness specified depending on the sequencing and other parameters as default. The resulting counts were imported into a DESeqDataSet object using the DESeq2 package [185] with a design formula of ~Group. Size factors were estimated and normalized counts were extracted and saved. Differential expression analysis was performed using DESeq with the created DESeqDataSet object, contrasted by sample groups, cooksCutoff and independentFiltering disabled, and otherwise default parameters.

### TMPRSS2 western blot

Cell lysates were prepared in RIPA buffer (150 mM NaCl, 1% NP-40, 0.5% DOC, 0.1% SDS, 50 mM Tris-HCl [pH 7.4] with addition of Halt^TM^ Protease and Phosphatase Inhibitor Cocktail [ThermoFisher: 78440]) and incubated on ice for 30 min. Lysates were clarified by centrifugation at 16,000 x g for 10 min at 4°C. Protein concentration was determined by bicinchoninic (BCA) protein assay (Pierce BCA Protein Assay Kit, ThermoFisher Scientific: 23227) and samples were resolved on NuPAGE 4%–12% Bis-Tris gels (Invitrogen) followed by transfer onto 0.4 µm nitrocellulose membranes. Membranes were incubated in blocking buffer (TBS + 5% milk) and incubated with primary antibody prepared in blocking buffer with 0.1% Tween-20. Membranes were washed 3X with TBS + 0.1% Tween-20 and incubated with LI-COR (Lincoln, NE, USA) IRDye® 680RD or IRDye® 800CW secondary antibodies. Membranes were imaged using an Azure (Dublin, CA, USA) 600 imaging system.

### Unbiased arrayed CRISPR KO screening

#### Screen overview

The content of each gRNA 384-well plate constituting the whole-genome library (61 library plates total) was transfected to 16 assay 384-well plates (976 assay plates total). Positive and negative control gene gRNAs were incorporated into vacant wells of each assay plate as described below. Huh-7.5-Cas9 cells were subsequently seeded into these assay plates. The 16 assay plates served as replicates for three distinct experimental conditions: 4 replicates for mock treatment followed by mock infection, 5 replicates for IFN-α2a treatment followed by SARS-CoV-2 infection, and 7 replicates for mock treatment followed by SARS-CoV-2 infection. Each day, three library 384-well plates were processed, along with their corresponding 48 assay plates. The full gRNA library, distributed across 61 384-well plates, was completed over a span of 21 days. For each set of plates, cell seeding was conducted on day 0, IFN-α2a treatment on day 4, SARS-CoV-2 infection on day 5, and cell fixation on day 6.

#### gRNA library preparation

A 0.1 nmol Edit-R Human Whole Genome crRNA Library (Horizon, cat. GP-005005-01) containing four crRNAs per gene and one gene per well (total 0.1 nmol crRNA/well) was resuspended in 80 µL of a 1.25 µM tracrRNA (Horizon, cat. U-002005-1000) 10 mM Tris-HCL pH 7.4 solution to create a 1.25 µM gRNA solution. The library was then aliquoted in 10 mM Tris-HCL pH 7.4 in several 96-well plate and 384-well plate copies using a Tecan Freedom EVO liquid handler. A single-use library copy containing a 40 µL/well of a 312.5 nM gRNA solution in the 384-well plate format was used in this study.

#### gRNA reverse transfection (day 0)

In each well of the 384-well assay plates, 40 µL of a transfection solution was prepared by combining 2% DharmaFect-4 transfection reagent (Horizon, cat. GP-T-2004-07A) in Opti-MEM (Gibco, cat. 31985070). This was added to 40 µL of a 312.5 nM gRNA library using a Thermofisher Multidrop Reagent Dispenser, yielding an 80 µL/well transfection mixture. The mixture was left to incubate at room temperature for 20 minutes. Simultaneously, assay plates were preloaded with 11 µL/well of serum-free media, which was formulated from DMEM, 1X Antibiotic-Antimycotic solution (Gibco, cat. 15240-062), and 1X NEAA, dispensed via a Thermofisher Multidrop Reagent Dispenser. Subsequently, 4 µL/well of the transfection mixture was dispensed into each of the assay plates (16 assay plates per library plate) using a Tecan Freedom EVO liquid handler. During this time, Huh-7.5 cells were prepared in media containing 25% FBS, 1X Antibiotic-Antimycotic solution, and 1X NEAA. A volume of 10 µL cells/well was added to the assay plates, again using a Thermofisher Multidrop Reagent Dispenser. Ultimately, each well contained 1,250 cells in a 25 µL final volume, with a composition of 25 nM gRNA, 10% FBS, 0.8X Antibiotic-Antimycotic, and 0.8X NEAA. Plates were then span at 200 g for 5 minutes. To minimize evaporation, plates were sealed with Breathe-Easy sealing membranes (Sigma-Aldrich, cat. Z380059) and placed in humid chambers constructed from a 245 mm x 245 mm dish containing a paper towel moistened with 15 mL of 1X Antibiotic-Antimycotic solution. Four assay plates were placed in each humid chamber and incubated at 37°C.

#### IFN-α2a treatment (day 1)

Each well received 5 µL of IFN-α2a (PBL, cat. 11101-2) in media (DMEM, 20% FBS, 1X Antibiotic-Antimycotic solution, 1X NEAA), using a Thermofisher Multidrop Reagent Dispenser, for a final concentration of 1 pM IFN-α2a in a final volume of 30 µL. Plates were then span at 200 g for 5 minutes and incubated at 37°C.

#### SARS-CoV-2 infection (day 5)

Each well received 212.5 PFU SARS-CoV-2 virus (titer determined on Vero E6 cells, see Virus Stocks section above) diluted in 5 µL of media (DMEM, 20% FBS, 1X Antibiotic-Antimycotic solution, 1X NEAA) for a final volume of 35 µL in the BSL3. Plates were then span at 200 g for 5 minutes and incubated at 37°C.

#### Fixing (day 6)

Each well received 50 µL of 20% neutral buffered formalin (Azer Scientific, cat. 20NBF-4-G) and plates were incubated overnight. The formalin mixture was then removed and each well received 50 µL of PBS.

#### IF staining

For IF staining of SARS-CoV-2 infected cells in the arrayed CRISPR KO screen (**Figs 1D-F**), as well as some focused experiments (**Fig 4**, **Fig 5G-I**, **Figs 7 C and E**, **Supp Fig 8**, **Supp Fig 9F**): the following solutions were prepared for both 96-well plate (96-wp) and 384-well plate (384-wp): PBS (Phosphate Buffered Saline), Perm Solution: Comprised of PBS with an added concentration of 0.1% Triton X100, Blocking Solution: PBS was mixed with 1% BSA. This solution was prepared a day in advance and filtered before use, PBST: PBS with 0.1% of Tween 20, Primary Antibody Solution: Genetex anti SARS-CoV-2 N poly rabbit antibody (GTX135357) at a dilution of 1:3000, Secondary Antibody Solution: AF647 anti-rabbit antibodies at a dilution of 1:3000 and Hoechst 33342 (10 mg/ml) at 1:10,000. Plates were stained on a Biotek EL406 Microplate Washer Dispenser using the following steps: 1. Priming: The washer was primed with 200 ml of each buffer: PBS, Perm Solution, Blocking Solution, and PBST. 2. First Washing Phase: Contents of the plates were aspirated. Plates were then washed with 50 µL/well (384-wp) or 200 µL/well (96-wp) of Perm Solution, followed by a slow shake for 3 seconds. 3. Permeabilization: A delay of approximately 1 minute was implemented for permeabilization, in addition to the time required to process all the plates (around 1 minute per plate). 4. Second Washing Phase: Plates were washed with 50 µL/well (384-wp) or 200 µL/well (96-wp) of PBS. Subsequently, 50 µL/well (384-wp) or 200 µL/well (96-wp) of Blocking Solution was added to the plates, followed by a slow shake for 3 seconds. 5. Blocking, autoclean, and Primary Antibody Priming: The washer was set to undergo an autoclean cycle with PBS for 30 minutes. Simultaneously, the syringe containing the Primary Antibody Solution was primed with 16 ml. 6. Third Washing Phase and First Antibody Dispensing: After aspirating the contents of the plates, 15 µL/well (384-wp) or 60 µL/well (96-wp) from the Primary Antibody Solution was added, followed by a slow shake for 3 seconds. 7. Primary Antibody Incubation, autoclean, and Secondary Antibody Priming: The washer was subjected to another autoclean cycle using PBS for 2 hours and 5 minutes. The syringe containing the Secondary Antibody Solution was primed with 16 ml during this period. 8. Fourth Washing Phase and Second Antibody Dispensing: Plates were washed with 50 µL/well (384-wp) or 200 µL/well (96-wp) of PBST, followed by a 2-second slow shake and aspiration. Then, 15 µL/well (384-wp) or 60 µL/well (96-wp) from the Secondary Antibody Solution was added, accompanied by a 3-second slow shake. 9. Secondary Antibody Incubation and autoclean: An autoclean cycle with PBS was initiated and lasted for 1 hour. 10. Final Washing Phase: Plates were washed with 50 µL/well (384-wp) or 200 µL/well (96-wp) of PBST. This was followed by two consecutive washes with 50 µL/well (384-wp) or 200 µL/well (96-wp) of PBS, incorporating a 2-second slow shake in each cycle. Finally, plates were left with 50 µL/well (384-wp) or 200 µL/well (96-wp) of PBS.

#### Imaging

Plates were imaged with a ImageXpress micro-XL and analyzed with MetaXpress (Molecular Devices).

#### Analysis

Analysis was conducted in R.

**Data Omission**: We excluded five library plates, constituting 8% of the total library, due to insufficient infection levels for accurate quantification.

**Normalization**: Two variables were subject to normalization—percentage of SARS-CoV-2 positive cells and the count of nuclei. The normalization steps were applied separately for the three screening conditions: mock treatment followed by mock infection, IFN-α2a treatment followed by SARS-CoV-2 infection, and mock treatment followed by SARS-CoV-2 infection. Data was first Z-scale normalized within assay plates:

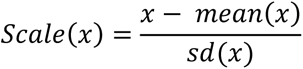

And then Z-scale normalized per row and per column to remove any spatial effects.

**Statistics:** a robust statistic accounting for technical and biological variability was applied using the below formula within the replicates of each gene:

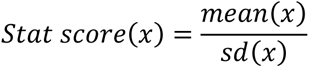

This statistic was further standardized by Z-scaling across all genes to produce our final z-

**Exclusion of genes influencing cell proliferation:** 224 genes with nuclei count z-score ≥ 2 and 388 genes with nuclei count z-score ≤ 2 in the mock treatment followed by mock infection condition were deemed to influence cell proliferation and excluded from subsequent analyses.

**Exclusion of genes not expressed in cell lines of interest (A549, Calu-3, Huh-7.5 cells) and in human lung cells.** Expression data from cell lines from [47, 48]. Expression data from tissues from [49]. Genes were considered expressed if they had at least one read count within exons.

### Gene set enrichment analysis

For the pathway analysis in **Supp Fig 2E**, we leveraged the FGSEA package [186] to perform Gene Set Enrichment Analysis (GSEA) using gene sets found in the Molecular Signatures Database (MSigDB) [187]: Reactome [188], KEGG [189], Wikipathways [190], Pathway Interaction Database [191], and Biocarta [192]. The analysis was conducted separately for two conditions: IFN-α pretreated SARS-CoV-2 infection and non-pretreated SARS-CoV-2 infection. We attributed a score to each pathway for both conditions:

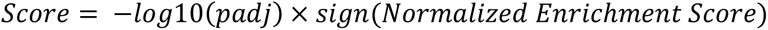

Each pathway was then attributed to one of nine quadrants (as in **Fig 1F**) based on its score in the IFN-α pretreated condition (axis x) versus non-pretreated condition (axis y), using padj ≤ 0.05 as a cutoff.

### Compilation of published large-scale omic studies on SARS-CoV-2

As a rule, we listed the genes classified as ‘hits’ by the authors of the respective studies. Below are some exceptions or clarifications:

#### Functional genetic screens

**Baggen, et al.** [80]: we used the “low stringency adjusted” analysis in Suppl Table 11. Proviral: p_value_neg ≤ 0.05 and log2 FC ≥ 1. Antiviral: p_value_pos ≤ 0.05 and log2 FC ≤ −1. We also used “High stringency” analysis in Suppl Table 7. Proviral: Gene is TMEM106B (log2 FC = 3.8 and p_value_neg = 0.08) or p_value_neg ≤ 0.05 and log2 FC ≥ 1 (no gene matched this criteria). Antiviral: p_value_pos ≤ 0.05 and log2 FC ≤ −1 (no gene matched this criteria). **Biering, et al.** [79]: in Supplementary Table 1, in Tab 1: LOF-enriched screen analysis, for proviral genes, we used FDR ≤ 0.05. In Tab 2: GOF-depleted screen analysis, proviral: FDR ≤ 0.05. In Tab 3: GOF-enriched screen analysis, for antiviral genes, we used FDR ≤ 0.05. **Chan, et al.** [78]: in Multimedia component 6, for Vero E6 (T16); UM-UC-4 (T23); HEK293+A+T (T12); HuH-7 (T15) and Calu-3 (T43), we considered gene as hits at FDR < 0.1 (as in Figure 4A). We listed the genes as proviral if differential ≥ 0 or antiviral if differential ≤ 0. **Daniloski, et al.** [77]: we used Table S1. FDR MOI1 ≤ 0.05 or FDR MOI3 ≤ 0.05. **Danziger, et al.** [11]: in S1 Table, we used the genes annotated as proviral or antiviral by the authors. **Gordon, et al.** [76]: for A549 +ACE2 in Table S6 or Caco2 in Table S7, for proviral genes, we used Averaged z-scores ≤ 2 and for antiviral genes, we used Averaged z-scores ≥ 2. **Grodzki, et al.** [75]: for VeroE6 in additional file 4, tab 4, we used FDR <0.25. For HEK293T +Cas9 Study 1 in additional file 6, tab17, we used FDR <0.25. For HEK293T +Cas9 Study 1 in additional file 7, tab7, we used FDR <0.25. **Hoffmann, et al.** [28]: for 37°C (Table S1E) and for 33°C (Table S1C), we selected proviral genes if FDR ≤ 0.05 and z-score ≥ 0 and antiviral genes if FDR ≤ 0.05 and z-score ≤ 0. **Hossain, et al.** [69]: in Figure 3E, we selected the top 15 genes in the spike-mNG axis by negative log robust rank aggregation. **Israeli, et al.** [74]: we used Supplementary Data 1. **Kaur, et al.** [15]: we used the genes labelled in Figure 1. **Le Pen et al. (this study):** we used z-score ≥ 2 for antiviral genes and ≤ 2 for proviral genes, see Unbiased arrayed CRISPR KO Screen analysis section above for more details. **Loo, et al.** [73]: we used the genes labelled in Figure 2. **Mac Kain, et al.** [18]: in Electronic Supplementary Material 5, for antiviral genes, we used the filter: pos|rank ≤13, for proviral genes, we used neg|rank ≤13. **Martin-Sancho, et al.** [12]: we used Table S3, “Lentivirus validated hits”. **Pahmeier, et al.** [72]: we used the genes labelled in Figure 6. **Rebendenne, et al.** [71]: for Calu3_Gattinara, we used for proviral genes: residual_z-score_avg ≥ 2.5 and for antiviral genes: residual_z-score_avg ≤ 2.5 (no gene). For VeroE6, proviral: residual_z-score_avg ≥ 2.5, and antiviral: residual_z-score_avg ≤ 2.5. For Caco2, proviral: residual_z-score_avg ≥ 2.5 and antiviral: residual_z-score_avg ≤ 2.5. For Calu3_Calabrese, proviral: residual_z-score_avg ≤ 2 and antiviral: residual_z-score_avg ≥ 2. **Rehfeld, et al.** [70]: In Table S1, we considered genes as hits for PRF-1 top eGFP-mCh or PRF-1 bottom eGFP-mCh if FDR ≤ 0.05. **Schneider, et al.** [31]: for both 37°C (Table_S1A) and 33°C (Table_S1B), we listed the gene as proviral if FDR ≤ 0.05 and z-score ≥ 0 and antiviral if FDR ≤ 0.05 and z-score ≤ 0. **Wang, et al.** [67]: we used Table S1. Proviral: Enrichment score ≤ 10^(−4). **Wei, et al. 2021** [66]: we used Table S1. For proviral genes, we used Cas9-v1 Avg. ≥ 2.5 & Cas9-v2 Avg. ≥ 2.5. Average between Cas9-v1 Avg. and Cas9-v2 Avg. is given in the table. For antiviral genes, we used Cas9-v1 Avg. ≤ −2.5 & Cas9-v2 Avg. ≤ −2.5. Average between Cas9-v1 Avg. and Cas9-v2 Avg. is given in the table. **Wei, et al. 2023** [65]: in Table S1, for Day 7 or Day 14, we used fdr ≤ 0.05 for positive regulators of ribosomal frameshifting or negative regulators of ribosomal frameshifting. **Wickenhagen, et al.** [13]: we used the genes labelled in Figure 1B. **Xu, et al.** [19]: in Huh-7.5 or A549-ACE2 cells, in untreated or in IFN-gamma treatment, we used Log10 p-value (mNG-High vs. mNG-Low Enrichment) ≥ 3. **Zhu et al.** [64]: In Supplementary Data 1, SARS-CoV-2 WT and 2 VOCs tested, for proviral genes, we used pos.score_wt ≤ 0.0005 or pos.score_alpha ≤ 0.0005 or pos.score_beta ≤ 0.0005.

#### Human genetic studies

**Degenhardt, et al.** [90]: we used Table 2 and added KANSL1 and TAC4, based on new analysis by Pairo-Castineira et al. 2023 [25]. **Kousathanas, et al.** [26]: we used Table 1 from Pairo-Castineira, et al. 2023 [25]. **Pairo-Castineira, et al. 2021** [98]: we used Table 1 from Pairo-Castineira, et al. 2023 [25]. **Pairo-Castineira, et al. 2023** [25]: we only considered variants near annotated genes (i.e., we excluded rs1073165). **Roberts, et al.** [83]: we used Figure 2. **Zhou, et al.** [95]: we used Table 1 and the p-values from COVID-19 hospitalization (European ancestry only).

#### SARS-CoV-2 protein interactomes

Davies, et al. [109], Laurent, et al. [102], Li, et al. [108], Samavarch-Tehrani, et al. [100], St-Germain, et al. [103], Stukalov, et al. [107]: we used the Supplementay Table 3 from [27]. Gordon, et al. [101]: we used the genes listed in Table S2. Liu, et al. [104] and May, et al. [105]: we used the genes listed in May, et al. [105] Table S5-new. Zhou, et al. [106]: we used the genes listed in Table S1. “SARS-CoV-2-human protein-protein interactions identified in this study.”

#### SARS-CoV-2 RNA interactomes

Flynn, et al. [114], Kamel, et al. [113], Labeau, et al. [112], Lee, et al. [115], Schmidt, et al. 2021 [111]: we used the Supplementay Table 3 from [27]. Schmidt, et al. 2023 [110]: we used Table S2, “Huh-7 interactome comparison tab”, genes listed in the following categories: “Huh-7 gRNA FDR5 HS” and “Huh-7 sgmRNA FDR5 HS”.

#### Altered phosphorylation states in SARS-CoV-2-infected cells

**Bouhaddou, et al.** [116]: for Vero E6 cells, we used Table S1, tab 1 “PhosphoDataFull” and filtered for adj.pvalue ≤ 0.05 & log2FC ≥ 1 or adj.pvalue ≤ 0.05 & log2FC ≤ −1 in at least three different time points.

### Generation of PLSCR1 KO cells

#### CRISPR KO

KO Huh-7.5 and A549-ACE2 cells were generated using two anti-PLSCR1 Edit-R crRNAs from Horizon Discovery (cat. CM-003729-02-0002 and CM-003729-04-0002) or non-targeting controls (cat. U-007501-01-05 and U-007502-01-05) resuspended with an equimolar amount of Edit-R tracrRNA (Horizon, cat. U-002005-20) to form sgRNAs. The sgRNAs were then co-transfected with Cas9-mKate2 mRNA (Horizon, cat. CAS12218) according to the manufacturer’s protocol. 24 to 48 hours after transfection, cells were examined for mKate2 signal and FACS sorted into bulk and single cell populations, gating on mKate2 signal. Bulk and single cell populations were then assessed for PLSCR1 expression by western blot to confirm KO.

#### Amplicon sequencing

Genomic DNA was isolated from a frozen cell pellet using the Qiagen DNeasy kit (Qiagen, cat. 69504) and treated with RNAse A in the optional RNA digestion step. The region of interest was then amplified using Q5 2x mastermix (New England Biolabs, cat. M0492S), 500 ng of template DNA, 0.5 µM of forward and reverse primers, and the following PCR conditions: 98°C for 30 seconds, followed by 98°C for 5 seconds, 64°C for 15 seconds, and 72°C for 20 seconds, repeating those steps 30 times before holding at 72°C for 2 minutes. The primers used when amplifying PLSCR1 genomic DNA from WT and KO Huh-7.5 and A549+ACE2 cells were RU-O-32687 (5’ AACATAGAGGTGATTATGATTTCGTCT) and RU-O-32526 (5’ GGAGGAGCTTGGATTTCTATCTAC). PCR reactions were run on a 1% agarose gel to confirm amplification. Amplicons were purified with a Zymo DNA clean and concentrator kit (Zymo, cat.

#### Western Blots

Cell pellets were collected and lysed in RIPA buffer (Thermo, catalog number 89900) with 1x Halt protease inhibitor cocktail and 1x EDTA (Thermo, catalog number 87786). Cell lysates were spun down in a refrigerated centrifuge at 15,000 g at 4°C for 15 minutes to pellet any cell debris, and the supernatant was collected and transferred to another tube. The collected samples were quantified by BCA assay (Thermo Scientific, cat. #23225). Before loading into the gel, we added sample buffer (Thermo, catalog number NP0007) with β-mercaptoethanol and heated the sample at 95°C for 10 minutes. Samples were allowed to cool back to room temperature before loading into 12% Bis-Tris 1.0 mm gels (Invitrogen, cat. #NP0321BOX). Proteins were electrophoretically transferred onto nitrocellulose membranes. Membranes were blocked with 5% fat-free milk in 1X TBS (Thermo, catalog number NP00061) and then incubated with primary antibody at 4°C overnight in 5% fat-free milk in 1x TBS with 0.5% Tween-20 (TBST). Primary antibody: rabbit anti-PLSCR1 polyclonal antibody (Proteintech, cat. #11582-1-AP) and mouse anti-β-actin antibody (Millipore Sigma, cat. A5316-100UL) as a loading control. After incubation, membranes were washed three times with 1x TBST and then incubated with fluorescently conjugated secondary antibodies for 2 hours at room temperature. Secondary antibodies: LI-COR IRDye goat anti-rabbit 800 and goat anti-mouse 680 (LI-COR cat. 926-32211 and 926-68070, respectively). Membranes were washed three times with 1X TBST, once with 1X TBS, then imaged on an Azure 600. For the western blot in Supplementary Figure 4, this protocol was modified slightly: proteins were electrophoretically transferred onto 0.22 µm polyvinylidene difluoride (PVDF) membranes, incubated with a primary antibody solution of rabbit anti-PLSCR1 polyclonal antibody (Proteintech, cat. #11582-1-AP) and polyclonal rabbit anti-RPS11 antibody (Abcam, cat. ab157101), a secondary antibody solution of goat anti-rabbit HRP (Invitrogen, cat. 31462) and visualized using a SuperSignal West Femto Maximum Sensitivity Substate kit (Thermo, cat. #34096).

### Cell viability assay

4,000 A549-ACE2 cells/well or 8,000 Huh-7.5 cells/well were seeded on day 0 in 100 µL media (DMEM, 10% FBS, 1X NEAA, 1X Penicillin-Streptomycin) in a 96-well plate. The next day, blasticidin selection was added as indicated in the figure to serve as a control for reduced cell viability. On day 4, cell viability was assessed by resazurin assay (Abcam, cat. ab129732) according to the manufacturer’s protocol.

### JAK-STAT inhibitor treatment

InSolution (Millipore, cat. 420097-500UG) was used according to the manufacturer’s instructions.

### Titration of IFN-a2a in CHIKV-infected cells

6,000 Huh-7.5 cells/well were seeded in 100 µL media (DMEM, 10% FBS, 1X NEAA). The following day, we treated cells with one of twelve concentrations of IFN-a2a (PBL, cat. 11101-2): 64 pM, 32 pM, 16 pM, 8 pM, 4 pM, 2 pM, 1 pM, 0.5 pM, 0.25 pM, 0.125 pM, 0.0625 pM, and 0 pM. The following day, the cells were infected with 2 µL of CHIKV-181/25-mKate2 (approximately 17,000 FFU per well, titer determined on Huh-7.5 cells) and fixed after 12 hours. Plates were stained with a 1:1000 dilution of Hoechst for at least 10 minutes before washing with PBS and imaging for mKate2 signal.

### RT-qPCRs on ISGs

Huh-7.5 and A549-ACE2 cells were seeded at densities of 36,000 or 18,000 cells/well, respectively, in 500 µL of media (DMEM, 10% FBS, 1X NEAA) in 24-well plates. The following day, a dilution series of IFN-α2a (PBL, cat. 11101-2) or IFN-β (PBL, cat. #11415) was prepared (64 pM, 32 pM, 16 pM, 8 pM, 4 pM, 2 pM, 1 pM, 0.5 pM, 0.25 pM, 0.125 pM, 0.0625 pM, and 0 pM) and 50 µL of each dilution added to the cells in duplicate. After 24 hours, the media was removed and the cells were washed with 1 mL of ice-cold PBS. 200 µL of RNA Lysis Buffer (Zymo Research, cat. #R1060-1-100) was added to the cells, and the plates were frozen at −20°C before RNA isolation.

RNA was extracted using the Zymo Quick RNA 96-kit (Zymo Research, cat. R1052) including DNaseI treatment, followed by cDNA synthesis using the SuperScript™ IV VILO™ Master Mix (Invitrogen, cat. 11756050) according to manufacturers’ instructions. qPCRs were conducted on a QuantStudio 3 cycler using the Taqman Fast Advance master mix (Life Technologies Corporation, cat. 4444965) and the following assays: *RPS11* (ThermoFisher 4331182; Hs01574200_gH), *IFI6* (ThermoFisher 4331182; Hs00242571_m1), *OAS1* (ThermoFisher 4331182; Hs00973635_m1). *IFI6* and *OAS1* were normalized to *RPS11* mRNA levels using the deltaCt method [193].

### PLSCR1 subcellular localization

#### IF staining

A549-ACE2 cells were plated onto #1.5, 12mm glass coverslips (Fisher Scientific, cat. #1254581) placed at the bottom of the wells of a 24-well plate. When confluent, the cells were fixed, permeabilized with 1% Triton X-100 for 5 minutes and blocked for 1 hour at room temperature with 1 mL of PBS-BGT (1x PBS with 0.5% bovine serum albumin, 0.1% glycine, 0.05% Tween 20). Afterward, the cells were incubated in a 1:500 dilution of 4D2 mouse anti-PLSCR1 antibody (Millipore Sigma, cat. #MABS483) in PBS-BG (1x PBS with 0.5% bovine serum albumin and 0.1% glycine) overnight at 4C with rocking. The cells were then washed twice with PBS-BGT before incubation with a secondary antibody solution of 1:1000 anti-mouse 588 (ThermoFisher, cat. #A-11001) and 1:1000 Hoechst dye (ThermoFisher, cat. #62249) in PBS-BG for two hours at room temperature, followed by three washes with PBS-BGT. For those images where cells were treated with IFN shown in **Supp Fig 11B**, A549-ACE2 cells were treated with 20 pM of IFN-α2a (PBL, cat. 11101-2) for 48 hours before harvest, then fixed and stained as described above. For those images where cells were infected with SARS-CoV-2 shown in **Supp Fig 11C**, A549-ACE2 cells were infected with approximately 3,400,000 PFU of SARS-CoV-2 (titer determined on Vero E6 cells) diluted in 200 µL Opti-MEM to achieve almost 100% infection. After 24 hours, cells were fixed and stained as described above, with slight modification to the antibody solutions. Primary antibody solution: 1:500 4D2 mouse anti-PLSCR1 and 1:3000 rabbit anti-SARS-CoV-2 N (Genetex, cat. GTX135357) in PBS-BG. Secondary antibody solution: 1:1000 goat anti-mouse 594, 1:1000 goat anti-rabbit 647, and 1:1000 Hoechst dye.

#### Imaging

The coverslips were mounted onto slides (Fisher Scientific, cat. #1255015) with Invitrogen ProLong Gold Antifade Mountant (Fisher Scientific, cat. # P36930). The slides were allowed to cure for 24 hours before the edges of the coverslips were sealed. For **Fig 5A**, the cells were imaged by confocal microscopy. Confocal images were acquired using Zeiss Zen Blue (v3.5) software on a LSM 980 point scanning confocal microscope (Zeiss) hooked to a Axio Observer.Z1 / 7 stand equipped with C Plan-Apochromat 63X/1.40 oil (RI:1.518 at 23°C) objective lens (Zeiss). CW excitation laser lines 405 nm and 488 nm were used to excite the fluorescence of DAPI and AF488 labeled samples. Emitted fluorescence were spectrally grated (410-483 nm for DAPI, 499-552 nm for AF488) to avoid fluorescence bleed through and were detected in MA-PMT (DAPI), and GaAsP-PMT (AF488). The confocal pinhole was set to 1AU for AF488, and the detector master gains were set within the linear range of detection (550-750V). For the IFN-treated and SARS-CoV-2 infected cells shown in **Supp Fig 11B,C**, these were mounted as described previously but imaged via widefield imaging. Widefield imaging was performed in the Rockefeller University’s Bio-Imaging Resource Center, RRID:SCR_017791, on a DeltaVision Image Restoration Microscope (Applied Precision, now Leica Microsystems) using inverted IX-70 stand (Olympus, Evident Scientific) and a 60X NA 1.42 oil immersion objective (Olympus), and filter sets for DAPI (Ex: 390 ± 18, Em: 435 ± 48), Alexa Fluor 594 (Ex: 575 ± 25, Em: 632 ± 60), Alexa Fluor 647 (Ex: 575 ± 25, Em: 676 ± 34). Images were acquired on a pco.edge sCMOS camera, and were subsequently deconvolved using the Deltavision image acquisition software (SoftWorx). Scanned images for both confocal and widefield imaging were saved as .czi files.

### Focus-forming assay on SARS-CoV-2-infected cells

In **Fig 5B**, Huh-7.5 and A549-ACE2 cells were cultured in media (DMEM with 5% FBS) and seeded at densities of 2 × 10^5^ and 1 × 10^5^ cells per well, respectively, in collagen-coated 12-well plates to reach 80-90% confluency by the day of infection. A 1:10 serial dilution of virus stock was made in Opti-MEM in five separate tubes. Media was aspirated from the cells, and the wells were washed with 1 ml of PBS before adding 200 µL of each virus dilution to the cells in triplicate. Plates were incubated at 37°C with 5% CO2 for 1 hour, rocking every 15 minutes for even virus distribution. A 1% methylcellulose overlay medium was prepared and mixed with complete growth media at 37°C; 2 ml of this overlay was added to each well after removing the virus inoculum. Plates were then incubated at 37°C with 5% CO2 for 48 for Huh-7.5 cells or 72 hours for A549-ACE2 cells. Cells were then fixed in final 10% neutral buffered formalin and IF stained as described in the Unbiased Arrayed CRISPR screen section. PLSCR1 KO and WT cells were compared at similar virus dilutions.

In **Fig 6G**, the above protocol was followed to titer SARS-CoV-2 strains on Huh-7.5 PLSCR1 KO cells. Then, Huh-7.5 WT and KO cells were seeded at 2 × 10^5^ cells per well in 1 mL of media (DMEM, 10% FBS, 1X NEAA) in 12-well plates to reach 80-90% confluency the next day. Media was aspirated from the cells, the wells were washed with 1mL of PBS, and then the cells were infected with approximately 50 FFU of SARS-CoV-2 (for each strain) as determined on PLSCR1 KO Huh-7.5 cells diluted in 200 µL of Opti-MEM. Plates were then incubated, overlayed with methylcellulose, fixed, and stained as described above.

### Transfection with SARS-CoV-2 replicon system

The SARS-CoV-2 replicon and the method for electroporation has been described previously [119]. Briefly, 6 × 10^6^ Huh-7.5 WT and PLSCR1 KO cells were electroporated at 710 V with 2 µg of SARS-CoV-2 N mRNA and 5 µg of replicon RNA. The cells rested for 10 minutes at room temperature before resuspending to a concentration of 300,000 cells/mL and plating 100 µL of cells into each well of a 96-well plate. After 24 hours, supernatant was collected from the replicon-transfected cells and assayed for *Renilla* luciferase activity according to kit instructions (Promega, cat. E2810).

### Transduction with SARS-CoV-2 spike/VSV-G-pseudotyped, single-cycle, replication-defective HIV-1 viruses

#### Virus preparation

SARS-CoV-2 spike/VSV-G-pseudotyped, single-cycle, replication-defective HIV-1 viruses (pCCNanoLuc/GFP) were prepared as in [120]. Plasmids were a kind gift of Theodora Hatziioannou and Paul D. Bieniasz (The Rockefeller University, NY, USA) [120, 194, 195]. One day before the transfection, 4 × 10^6^ 293T cells were seeded in a 10 cm dish. One hour prior to transfection, the growth media in the dish was replaced with 9 mL of fresh media containing 2% serum. A 1,000 µL transfection mixture was prepared, comprising the diluent (a 150 mM NaCl solution prepared with sterile cell culture water), 5 µg of HIV GP plasmid, 5 µg of pCLG plasmid, and either 2.5 µg of SARS-CoV-2 spike Δ19 or 1 µg of pHCMV.G plasmid, ensuring the total plasmid content did not exceed 12.5 µg. After brief vortexing, 50 µL of PEI (1 mg/mL, Polysciences cat. 23966) was added to achieve a 1:4 DNA/PEI ratio. The mixture was vortexed for 5 seconds and then allowed to sit for 20 minutes in a hooded environment. Following gentle mixing by pipetting, 1 mL of the transfection mixture was added to the 10 cm dish. Media was changed 12 hours post-transfection, and the supernatant was harvested and filtered through a 0.2-micron filter 48 hours post-transfection, then stored at −80°C.

#### Transduction of PLSCR1 KO or WT A549-ACE2 cells

Seeded in 96-well plates, 6,000 A549-ACE2 cells per well were cultured in 100 µL of media. After two days, either 10 µL of SARS-CoV-2 spike pseudotyped virus or 0.01 µL of VSV-G pseudotyped virus were diluted in a final volume of 100 µL of media and added to the wells to yield comparable NanoLuc signals. Plates were then spun at 200 g for 5 minutes and incubated at 37°C. Two days post-infection, the media was aspirated and replaced with 50 µL of NanoGlo solution, sourced from the Promega Nano-Glo Luciferase Assay kit (Promega, N1110), with a substrate to buffer ratio of 1:100. NanoLuc signal was subsequently quantified using a Fluostar Omega plate reader.

#### Transduction of siRNA-treated HEK293T cells

Seeded in 96-well plates, 1,600 HEK293T, HEK293T-ACE2, or HEK293T-ACE2-TMPRSS2 cells were cultured in 80 µL of media. The next day, a 20 µL transfection mixture made of Opti-MEM, 1% DharmaFECT1 (Horizon, T-2001-03), and 250 nM siRNA, PLSCR1 ON-TARGETplus SMARTpool siRNA (Horizon, cat. L-003729-00-0005) or non-targeting control (Horizon, cat. D-001810-10-05) was added to the cells. The final concentration of siRNAs was 25 nM. After two days, either 2 µL of SARS-CoV-2 spike pseudotyped virus or 0.2 µL of VSV-G pseudotyped virus were diluted in a final volume of 100 µL of media and added to the wells to yield comparable NanoLuc signals. Plates were then span at 200 g for 5 minutes and incubated at 37°C. Two days after infection, NanoLuc signal was quantified as described in the “Infection of PLSCR1 KO or WT cells” section.

### Infection of siRNA-treated A549 cells with SARS-CoV-2

Seeded in 96-well plates, 1,000 A549, A549-ACE2, or A549-ACE2-TMPRSS2 cells were cultured in 80 µL of media. On the same day, a 20 µL transfection mixture made of Opti-MEM, 1% DharmaFECT1 (Horizon, T-2001-03), and 250 nM siRNA, PLSCR1 ON-TARGETplus SMARTpool siRNA (Horizon, cat. L-003729-00-0005) or non-targeting control (Horizon, cat. D-001810-10-05) was added to the cells. The final concentration of siRNAs was 25 nM. Three days after transfection, the cells were infected by adding 34,000 PFU of SARS-CoV-2 (titer determined on Vero E6 cells) diluted in 10 µL media to each well. Plates were then spun at 200 g for 5 minutes and incubated at 37°C. Staining and readout as described above in the “Unbiased arrayed CRISPR KO screening” section.

### Pan-virus infection of PLSCR1 KO cells

A549-ACE2 cells were seeded at a density of 6,000 cells/well in 96-well plates in 90 µL media. The following day, 10 µL diluted virus was added to each well. Virus concentrations as follow: CHIKV-mKate, 0.05 µL virus stock per well (titer 8.5 × 10^6^ PFU/mL determined in Huh-7.5 cells); hCoV-NL63, 10 µL virus stock per well (titer 1.4 × 10^5^ PFU/mL); hCoV-OC43, 10 µL virus stock per well (titer 1.06 × 10^7^ PFU/mL); hPIV-GFP, 0.05 µL virus stock per well; HSV1-GFP, 0.5 µL virus stock per well (titer 2.4 × 10^8^ PFU/mL determined on Vero E6 cells); IAV WSN, 0.5 µL virus stock per well; SARS-CoV-2, 0.5 µL virus stock per well (titer 3.4 × 10^6^ PFU/mL determined on Vero E6 cells); SINV-Toto1101-mScarletI, 10 µL virus stock per well; VEEV-EGFP, 0.005 µL virus stock per well (titer 1.45 × 10^9^ PFU/mL determined on BHK-21); VSV-GFP, 0.05 µL virus stock per well; YFV_17D, 5 µL virus stock per well. The cells were fixed, stained and imaged as described in the *Unbiased arrayed CRISPR KO screen* section. Fluorescent viruses were not stained: the fluorescent signal was used as a reporter. We used the following primary antibodies when applicable: anti-dsRNA (J2) mouse (Nordic MUbio, cat. 10010200) diluted 1:500 was used for hCoV-NL63 and hCoV-OC43, anti-IAV mouse (Millipore, cat. MAB8257) diluted 1:3000, anti-YFV mouse (Santa Cruz Biotechnology, cat# sc-58083) diluted 1:500, anti-SARS2-S rabbit (Genetex, cat. GTX135357) diluted 1:3000.

Huh-7.5 cells were transfected with a 1:200 dilution of Dharmafect 4 (Horizon, cat. T-2004-01) and 25 nM ON-TARGETplus SMARTpool siRNAs (Horizon Discovery) in 96-well plates. The cells were infected three days after siRNA transfection with hCoV-NL63 or hCoV-OC43 or four days after siRNA transfection with SARS-CoV-2 or hCoV-229E. IF and imaging as described above.

### Measuring extracellular ACE2 expression using flow cytometry

A549-ACE2 cells were detached from the wells using Accutase (Innovative Cell Technologies, Inc., Cat: AT-104) for 5 minutes at 37°C. Following detachment, cells were rinsed once with PBS and stained with LIVE/DEAD Fixable Aqua (Invitrogen, Cat: L34966, diluted 1:1000 in PBS) for 15 minutes at room temperature in the dark. After staining, cells were rinsed once more with PBS. Then, cells were incubated with FcR blocking reagent (Miltenyi Biotec, 1:50) and ACE2-AF488 (Sino Biological, Cat: 10108-MM37-F-100, 1:30) in FACS buffer (2% FBS in PBS) for 30 min at 4°C in the dark. Following incubation, cells were washed twice with FACS buffer and subjected to flow cytometry analysis. Samples were acquired on an LSRII flow cytometer (BD Biosciences), and the results were analyzed with FlowJo software (Tree Star). Before analyzing cell surface ACE2 expression, gating procedures were implemented to exclude dead cells and doublets from the analysis.

### SARS-CoV-2 variant infections

For **Figure 6A-E**, Huh-7.5 cells were plated at 7,500 cells/well in 100 µL of media (DMEM, 10% FBS, 1X NEAA) in 96-well plates. The next day, cells were infected with quantities of virus that yielded comparable percent infections in the WT cells, then spun at 200 g for 5 minutes and incubated at 33°C. The quantities of virus used were as follows: 0.1 µL/well for parental, 0.05 µL/well for beta, 1 µL/well for delta, 0.5 µL/well for Omicron BA.5, and 0.05 µL/well for Omicron XBB.1.5. The infected cells were fixed after 24 hours and stained as described previously.

### Growth curves using the parental SARS-CoV-2 strain and Omicron (XBB.1.5)

Huh-7.5 WT and PLSCR1 KO cells were plated at 100,000 cells/well in 1mL media (DMEM, on PLSCR1 KO Huh-7.5 cells) with the New York 2020 isolate of SARS-CoV-2 (“parental”) and SARS-CoV-2 Omicron XBB.1.5. Plates were incubated for 1hr at 37°C. Inoculum was removed and saved as the 0hr timepoint. Cells were washed twice with PBS. Fresh media was added, then collected as the 1hr timepoint. Fresh media was added again, and the cells were incubated at 33°C. All supernatant was collected from the cells again at 24 hours, 48 hours, and 72 hours, and was replaced with fresh media. Collected supernatant was frozen at −80°C until titering. Supernatants were titered by plaque assay as follows: Vero E6 cells were plated at 200,000 cells/well in 12-well plates. The following day, collected supernatant was tenfold serially diluted, the media removed from the plates, and 300 µL diluted virus added. Each sample of supernatant was titered in duplicate. Cells were incubated with virus for 1 hour at 37°C, then inoculum was removed and 1mL of 1% methylcellulose (supplemented with 5% FBS). Plates were incubated for 3 days at 37°C, then fixed with 2mL of 7% formaldehyde and stained with crystal violet before being rinsed with water, left to dry overnight, and the plaques counted.

### Generating and infecting cells overexpressing ISGs (Including PLSCR1)

#### Plasmid sources

WT PLSCR1, IFITM2, and LY6E over-expression lentiviral plasmids originated from the SCRPSY library [196]. CH25H originated from the Thermo Scientific Precision LentiORF Collection. RRL.sin.cPPT.CMV/Flag-NCOA7 variant 6.IRES-puro.WPRE (CG494) was a gift from Caroline Goujon (Addgene plasmid # 139447) [161].

#### Plasmid cloning

N-terminal 3x FLAG-tagged PLSCR1, C-terminal 3x FLAG-tagged PLSCR1, and PLSCR1 H262Y in the SCRPSY vector, used in **Figs 4B and 7D**, were generated by designing and ordering large dsDNA gene blocks of PLSCR1 that contained the desired mutations from IDT. These gene blocks were cloned into the PLSCR1-SCRPSY vector [196] and confirmed by sequencing using Plasmidsaurus (see **Supp Table 18** for sequences).

PLSCR1 variants of interest (**Fig 7C and Supp Fig 12**) were cloned by site directed mutagenesis using the NEB Quick Protocol for Q5® Site-Directed Mutagenesis Kit (cat. E0552) according to the manufacturer’s instructions. For G20V (rs79551579), we used the primers JLP621-O-01/02; for A29T (rs41267859), we used the primers JLP621-O-03/04; for I110fs rs749938276, we used the primers JLP621-O-07/08; and for H262Y (rs343320), we used the primers JLP621-O-09/10. Primer sequences are listed in **Supp Table 20**.

#### Lentivirus production

Lentivirus were generated in Lenti-X 293T cells by transfecting 200 ng VSV-G plasmid, 700 ng Gag-Pol plasmid, and 1100 ng plasmid of interest with lipofectamine 2000 in DMEM supplemented with 5% FBS. Media was changed 4-6 hours later, and lentivirus harvested at 24 and 48 hours. Lentivirus-containing supernatants from both timepoints were pooled, then filtered through a 0.45 µM filter before aliquoting into 2 mL tubes and freezing at −80°C until use.

#### Cell transduction

0.3 million cells were transduced in suspension in a 12-well plate using a serial dilution of each lentivirus in DMEM (Fisher Scientific, cat. #11995065) supplemented with 5% FBS (HyClone Laboratories, Lot. #KTH31760) and 4 µg/mL polybrene (Millipore, cat. #TR-1003-G). The cells were then spinoculated at 37°C for 1 hour at 1000 x g. The following day, the cells were split into two 6-well plates. After 24 hours, one of the duplicates was treated with 2 µg/mL puromycin to select for transduced cells. Further experiments were carried out using the cells that had approximately 30% transduction before selection.

#### SARS-CoV-2 infection

In **Fig 7D**, Huh-7.5 and A549+ACE2 cells were plated at 6,000 cells/well and 3,000 cells/well, respectively, in 100 µL of media (DMEM, 10% FBS, 1X NEAA) in 96-well plates. The following day, cells were treated with IFN (10pM for Huh-7.5 cells and 20pM for A549+ACE2 cells). On the third day, the Huh-7.5 cells were infected by adding 10 µL of media containing 0.1 µL of virus (titer of 3.4 × 106 PFU/ml, as measured on Vero E6 cells) to each well and the A549+ACE2 cells were infected by adding 10 µL of media containing 1 µL of virus to each well. After adding the virus, the plates were spun at 200 g for 5 minutes and incubated at 37°C. Plates were harvested the next day by fixing and staining as described above

In **Fig 7C** and **Supp Figs 8A** and **12**, A549+ACE2 cells were plated at 7,000 cells/well in 100 µL of media (DMEM, 10% FBS, 1X NEAA) in 96-well plates. The following day, cells were infected by adding 10 µL of media containing 1 µL of virus (titer of 3.4 × 106 PFU/ml, as measured on Vero E6 cells) to each well as described above.

#### IF staining and imaging

In **Figs 7C-E and Supp Figs 8A and 12**, cells were stained for IF as described in the *Unbiased arrayed CRISPR KO screening* section, with different primary and secondary antibody solutions. The primary antibody solution was a 1:3000 dilution of rabbit anti-SARS-CoV-2 nucleocapsid polyclonal antibody (Genetex, cat. #GTX135357) and a 1:500 dilution of 4D2 mouse anti-PLSCR1 antibody (Millipore Sigma, cat. #MABS483) in PBS, and the secondary antibody solution was 1:1000 goat anti-rabbit 594 (ThermoFisher, cat. #A-11012) or 1:1000 goat anti-rabbit Alexa Fluor™v 647 (ThermoFisher, cat. **#**A-21245), 1:1000 goat anti-mouse Alexa Fluor™ 488 (Thermofisher, cat. #A-11001), and 1:1000 Hoechst dye (ThermoFisher, cat. #62249) in PBST.

### SARS-CoV-2 infection of human SV40-fibroblasts-ACE2

#### Patients

Written informed consent was obtained in the country of residence of the patients, in accordance with local regulations, and with institutional review board (IRB) approval. Experiments were conducted in the United States in accordance with local regulations and with the approval of the IRB. Approval was obtained from the Rockefeller University Institutional Review Board in New York, USA (protocol no. JCA-0700).

PLSCR1 H262Y was genotyped using the primers JLP616-O-1-4 listed in **Supp Table 20**.

#### Generation of human SV40-fibroblasts ACE2 stable cell lines

*ACE2* cDNA was inserted with In-Fusion cloning kit (Takara Bio) and using the XhoI and BamHI restriction sites into linearized pTRIP-SFFV-CD271-P2A in accordance with the manufacturers’ instructions. We checked the entire sequence of the *ACE2* cDNA in the plasmid by Sanger sequencing. Then, HEK293T cells were dispensed into a six-well plate at a density of 8 x10^5^ cells per well. On the next day, cells were transfected with pCMV-VSV-G (0.2 µg), pHXB2-env (0.2 µg; NIH-AIDS Reagent Program; 1069), psPAX2 (1 µg; Addgene plasmid no. 12260) and pTRIP-SFFV-CD271-P2A-ACE2 (1.6 µg) in Opti-MEM (Gibco; 300 µL) containing X-tremeGene-9 (Sigma Aldrich; 10 µL) according to the manufacturers’ protocol. After transfection for 6 h, the medium was replaced with 3 mL fresh culture medium, and the cells were incubated for a further 24 h for the production of lentiviral particles. The viral supernatant was collected and passed through a syringe filter with 0.2 μm pores (Pall) to remove debris. Protamine sulfate (Sigma; 10 µg/mL) was added to the supernatant, which was then used immediately or stored at −80 °C until use.

For the transduction of SV40-fibroblasts with ACE2, 5 x10^5^ cells per well were seeded in six-well plates. Viral supernatant was added (500 µL per well). The cells were then further incubated for 48 h at 37 °C. Cells were keep in culture and after 8 days, transduction efficiency was evaluated by CD271 surface staining with CD271 AlexaFluor 647, 1:200 dilution (BD Pharmigen 560326). MACS-sorting was performed with CD271 positive selection beads (Miltenyi Biotec) if the proportion of CD271-positive cells was below 80% [8, 177].

#### Infection

5,000 cells per well were seeded in a 96-well plate and infected the next day with SARS-CoV-2 at MOI = 0.05, using a titer determined on Vero E6 cells. Cells were fixed at 2 dpi, stained and imaged as described in the *Unbiased arrayed CRISPR KO screen* section.

## Acknowledgments

We thank Georgia McClain for reading and editing this manuscript. We thank the staff at the Laboratory of Virology and Infectious Disease: Ellen Castillo, Michela De Santis, Arnella Norris, Aileen O’Connell, Santa Maria Pecoraro Di Vittorio, Glen Santiago, and Sonia Shirley. Ching-Wen Chang and Lihong Liu (Columbia University, NY, USA), and Theodora Hatziioannou, Viren Baharani and Paul D. Bieniasz (The Rockefeller University, NY, USA) generously provided plasmids, instructions to generate SARS-CoV-2 spike-pseudotyped, single-cycle, replication-defective HIV-1 viruses [120, 194, 195], and assisted us with data interpretation. Oded Danziger and Brad R. Rosenberg (Department of Microbiology at the Icahn School of Medicine at Mount Sinai, NY, USA) kindly provided the A549-ACE2 cells [11] used in this study. FACS was conducted at the Flow Cytometry Resource Center at Rockefeller University. mRNA-seq was performed by the Genomics Resource Center at The Rockefeller University and by Novogene. Confocal microscopy was performed in the Rockefeller University’s Bio-Imaging Resource Center, RRID:SCR_017791. We thank Ankit Patel, Sales Manager at Horizon Discovery.

Trim Galore was developed at The Babraham Institute by @FelixKrueger, now part of Altos Labs. The Genotype-Tissue Expression (GTEx) Project [49] was supported by the Common Fund of the Office of the Director of the National Institutes of Health, and by NCI, NHGRI, NHLBI, NIDA, NIMH, and NINDS. The data used for the analyses described in this manuscript were obtained from the EMBL-European Bioinformatics Institute portal, “https://www.ebi.ac.uk/gxa/experiments/E-MTAB-5214/Results”, on October 1^st^, 2020.

Work in the Laboratory of Virology and Infectious Disease was supported by NIH grants P01AI138398-S1, 2U19AI111825, R01AI091707-10S1, and R01AI161444; a George Mason University Fast Grant; the G. Harold and Leila Y. Mathers Charitable Foundation; the Meyer Foundation; the Pilot Project Robertson Therapeutic Development Fund at The Rockefeller University; and the Bawd Foundation. The Laboratory of Human Genetics of Infectious Diseases is supported by the Howard Hughes Medical Institute, the St. Giles Foundation, NIH (R01AI163029, UL1TR001866), the French National Research Agency (ANR) under the “Investments for the Future” program (ANR-10-IAHU-01), the Integrative Biology of Emerging Infectious Diseases Laboratory of Excellence (ANR-10-LABX-62-IBEID), the French Foundation for Medical Research (FRM) (EQU201903007798), ANR GENVIR (ANR-20-CE93-003) project, the ANR-RHU COVIFERON Program (ANR-21-RHUS-08), the French Foundation for Medical Research (FRM) (EQU201903007798), the European Union’s Horizon 2020 research and innovation program under grant agreement No. 824110 (EASI-genomics), the HORIZON-HLTH-2021-DISEASE-04 program under grant agreement 01057100 (UNDINE), the Square Foundation, William E. Ford, General Atlantic’s Chairman and Chief Executive Officer, Gabriel Caillaux, General Atlantic’s Co-President, Managing Director and Head of Business at EMEA, and the General Atlantic Foundation. J.L.P. was supported by the Francois Wallace Monahan Postdoctoral Fellowship at The Rockefeller University and the European Molecular Biology Organization Long-Term Fellowship (ALTF 380-2018). G.P. was supported by the James H. Gilliam Fellowship for Advanced Study from the Howard Hughes Medical Institute and the Graduate Research Fellowship Program from the National Science Foundation (FAIN 1946429). M.B. was supported by a Swiss National Science Foundation fellowship (P500PB_203007).

## Financial Disclosure

The funders had no role in study design, data collection and analysis, decision to publish, or preparation of the manuscript.

## Data and code availability

mRNA-seq data is available on the Sequence Read Archive (SRA), hosted by the National Center for Biotechnology Information (NCBI), under BioProject PRJNA1138251 and BioSamples SAMN42689767-SAMN42689770 corresponding to **Supp Tables 2 and 3**, BioSamples SAMN42689771-SAMN42689774 corresponding to **Supp Tables 4 and 5**, BioSamples SAMN42689775-SAMN42689776 corresponding to **Supp Tables 14 and 15**, BioSamples SAMN42689777-SAMN42689778 corresponding to **Supp Tables 16 and 17**.

Code and Supporting Information are available on Dryad (DOI: 10.5061/dryad.6q573n65k).

## Supplementary Figure Legends

**Supplementary Figure 1.**
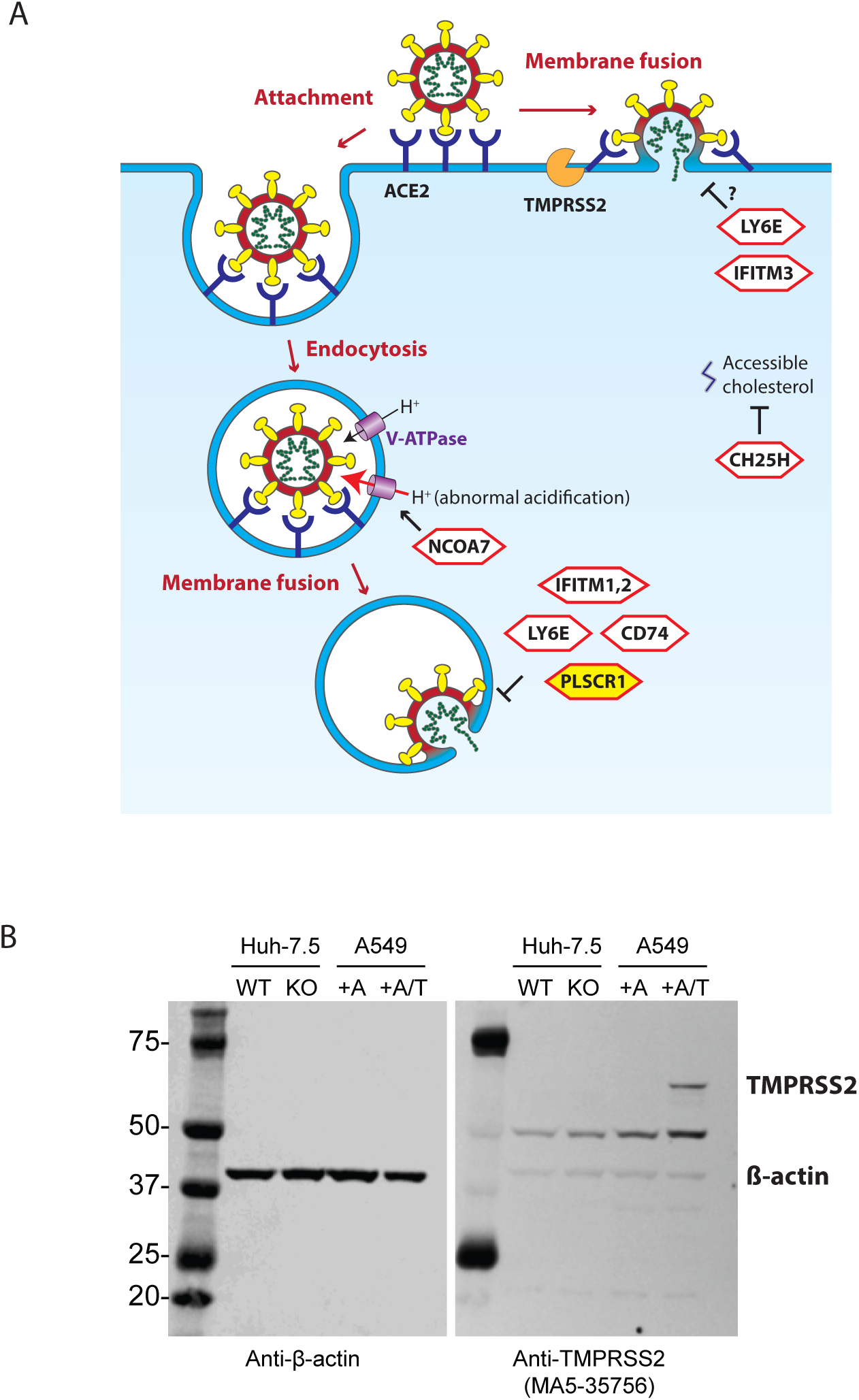
A. A schematic of SARS-CoV-2 entry and the sites where known ISG entry restriction factors function is shown, adapted from [1]. CD74 suppresses endolysosomal cathepsins, enzymes that process certain viral glycoproteins to make them fusion-competent [148, 198]. CH25H facilitates the sequestration of accessible cholesterol, which results in decreased virus-cell membrane fusion and viral entry [14, 63]. NCOA7 accelerates the acidification of the lysosome, leading to the degradation of viral antigens [161, 162]. The role of IFITM1-3 proteins in SARS-CoV-2 entry has been the subject of numerous studies, which have sometimes yielded varying conclusions [1, 122, 199–205]. However, it appears that (i) IFITM2 restricts WT SARS-CoV-2 endosomal entry [122], (ii) recent SARS-CoV-2 variants may evade IFITM2 [199], and (iii) IFITM3 KO mice are hyper-susceptible to WT SARS-CoV-2 [206]. LY6E and PLSCR1 restrict virus-cell membrane fusion at the endosome through unknown mechanisms, see [19, 121] and this study. B. TMPRSS2 western blot in Huh-7.5-Cas9 cells, both WT and TMPRSS2 KO, conducted as in the whole-genome arrayed screen (Fig 1), and in A549-ACE2 (A549 + A) and A549-ACE2-TMPRSS2 (A549 + A/T) cells.

**Supplementary Figure 2.**
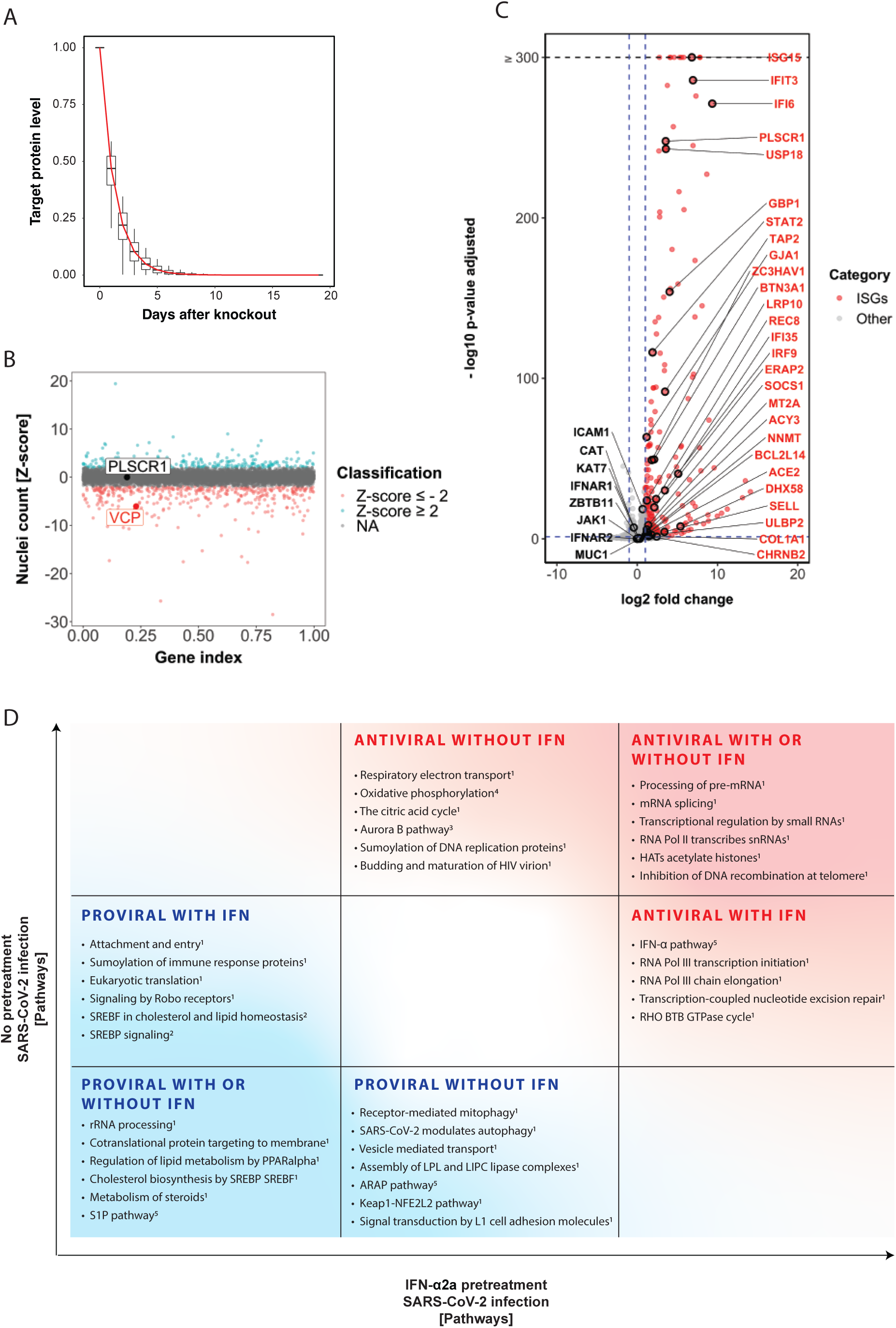
A. Model of protein levels of the gene product over time after complete KO, considering the Huh-7.5 cell division rate (1.7 times per day) and protein half-lives in hepatocytes from [207]. B. Nuclei count (z-score) in arrayed genetic screen without SARS-CoV-2 infection. Examples of VCP (essential gene) and PLSCR1 (SARS-CoV-2 antiviral hit) are plotted. C. Volcano plot of Huh-7.5 cells mRNA-seq treated with 0.5 nM IFN-α2a for 24 h as in **Fig 1B**. D. Gene set enrichment analysis conducted on the arrayed CRISPR KO screens data represented in figS 1 and 2. Description of the top pathways ranked by p-value for each quadrant. Databases: ^1^Reactome; ^2^WikiPathways; ^3^Pathway Interaction Database; ^4^KEGG; ^5^Biocarta. The data underlying this Figure can be found in **Supp Table 1**.

**Supplementary Figure 3.**
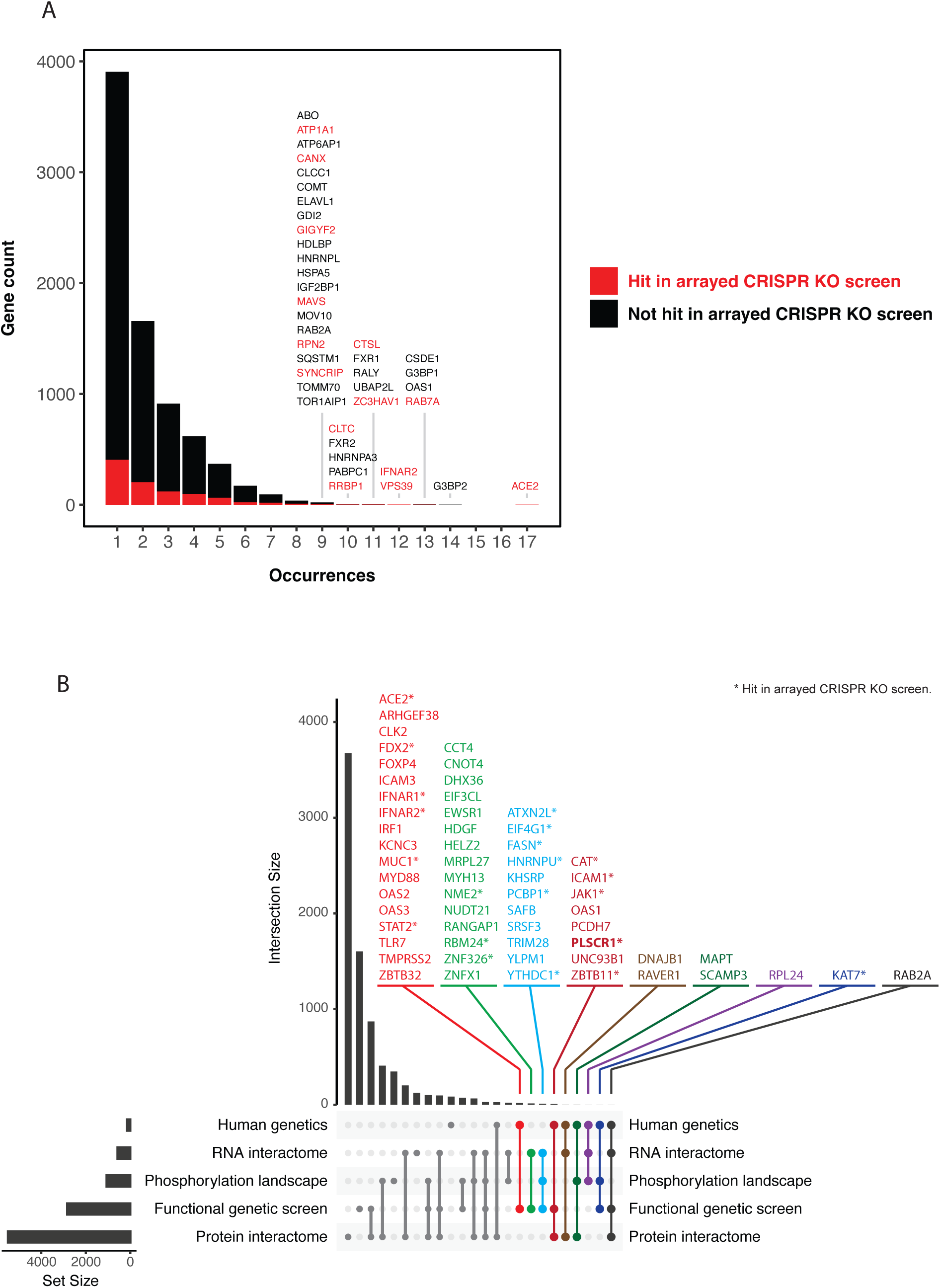
A. Occurrence of human genes interacting with SARS-CoV-2 drawn from a selection of 67 large-scale studies. The occurrence reflects the number of independent studies finding each gene as significant. B. Upset plot on data as in (A), showing the overlap in significant genes in large-scale SARS-CoV-2 studies by category.

**Supplementary Figure 4.**
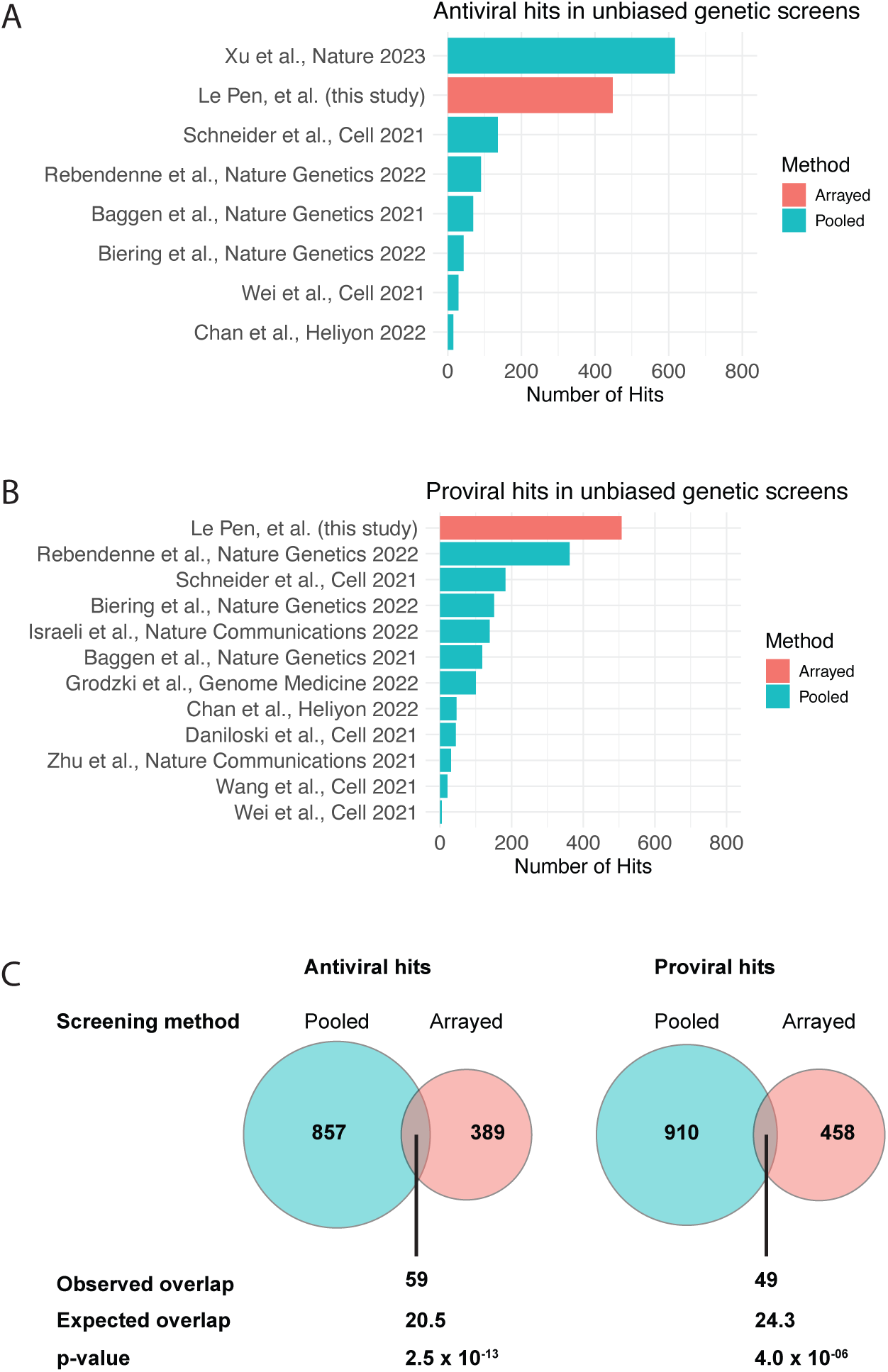
Cross comparison between proviral and antiviral hits reported in this study and fifteen published whole-genome pooled screens [19, 31, 64–67, 69–71, 74, 75, 77–80]. P-value: Fisher’s exact test.

**Supplementary Figure 5.**
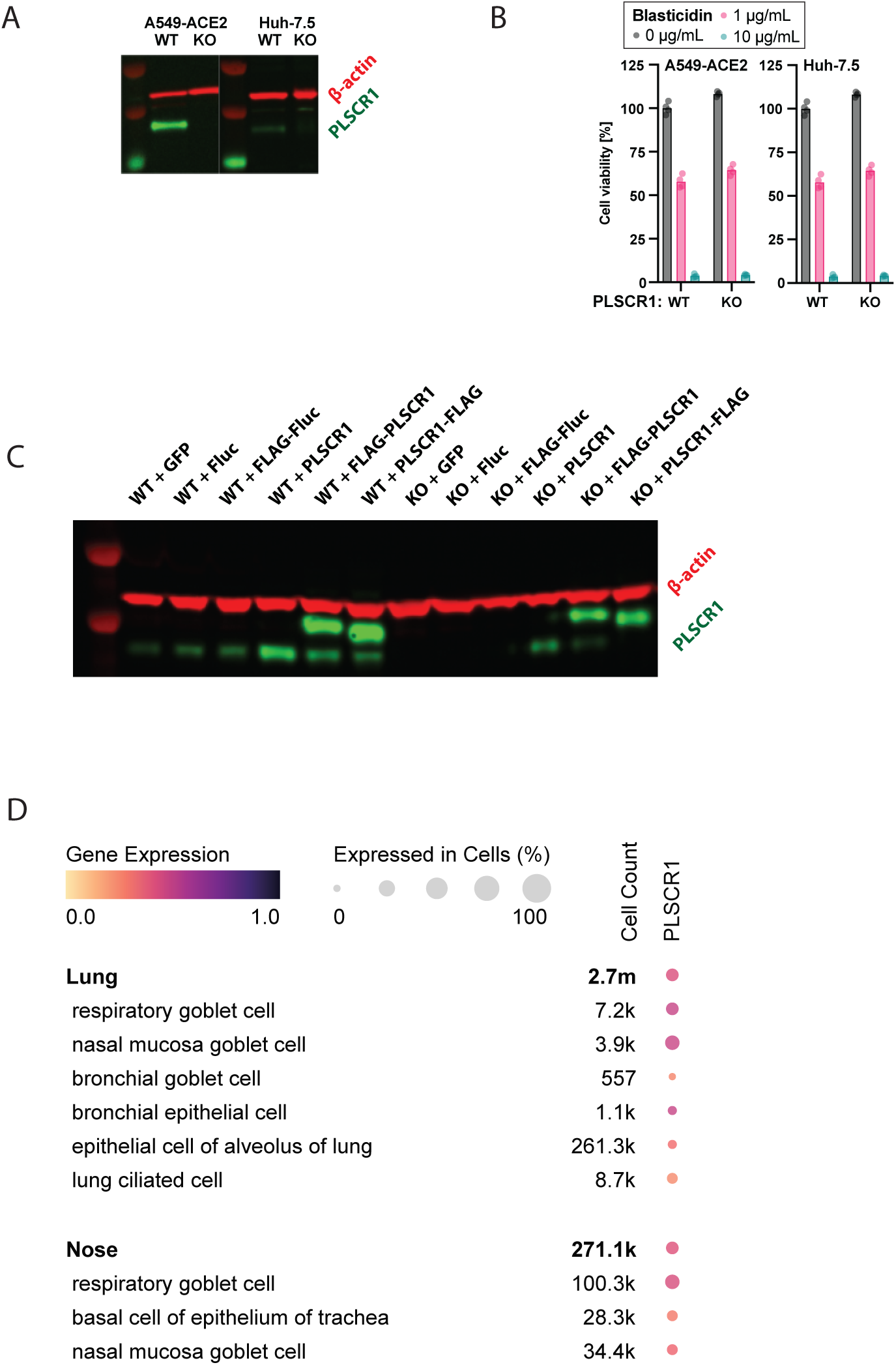
A. Western blot on PLSCR1 KO cells against PLSCR1 (green) and ß-actin (red). B. Cells as indicated were seeded at similar density, treated or not with Blasticidin (used here as a control to decrease cell viability), and cultured for 4 days before resazurin cell viability assay. n = 4 independent wells. C. Western blot on PLSCR1 WT and KO A549-ACE2 cells against PLSCR1 (green) and ß-actin (red). D. Single-cell RNA-seq data from [118], showing the *PLSCR1* mRNA expression in some SARS-CoV-2 target cells. The data underlying this Figure can be found in **Supp Table 1**.

**Supplementary Figure 6.**
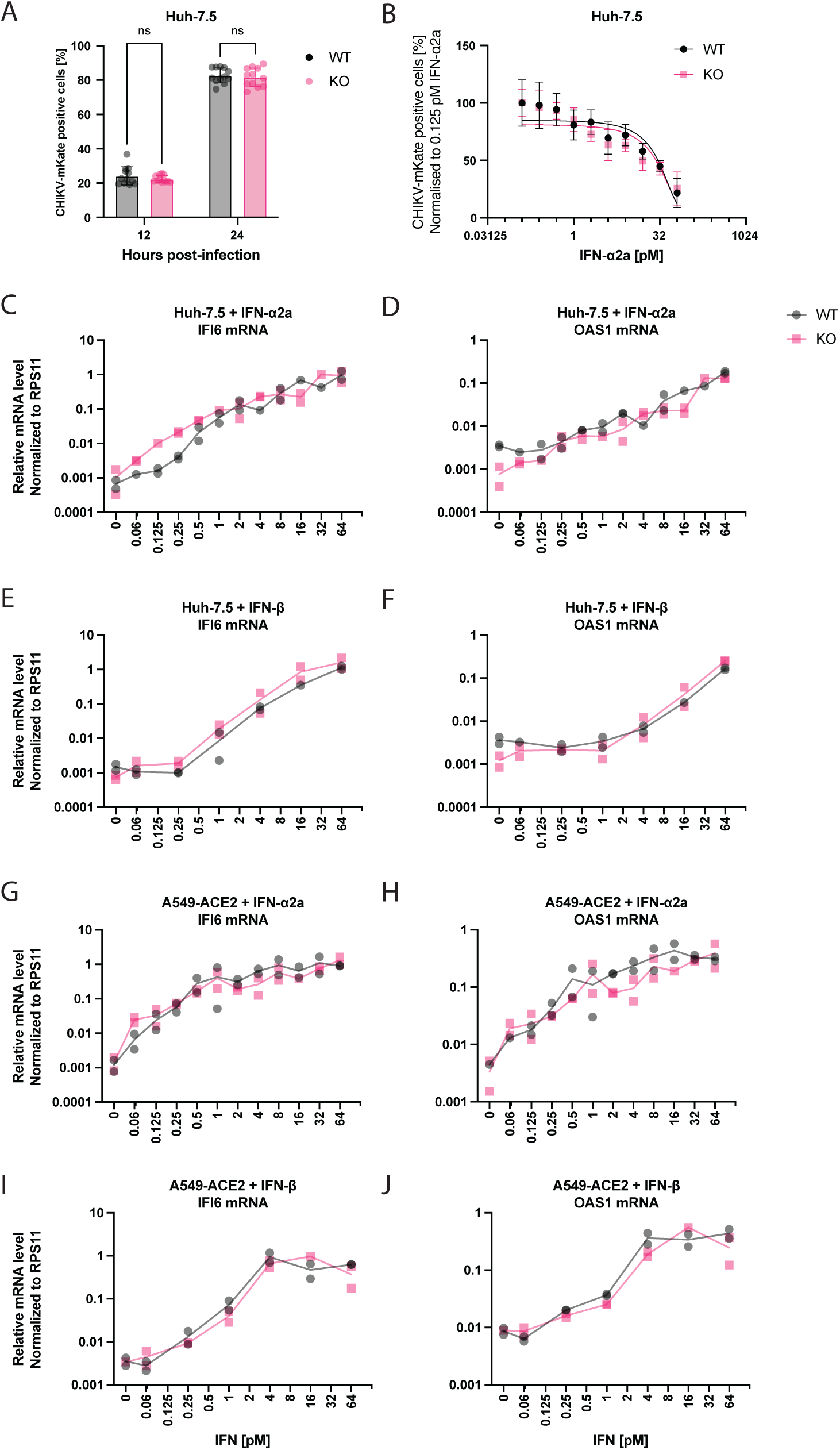
A. Huh-7.5 WT and PLSCR1 KO cells were infected with 17,000 FFU of CHIKV 181/25 mKate2 for 12 or 24 hours, then fixed, IF stained for nuclei, and the percentage of positive cells determined by imaging for mKate2 reporter signal. n = 12 separate wells infected on the same day. Error bars represent sd. ns, non-significant; two-way ANOVA. B. Huh-7.5 cells, PLSCR1 KO as indicated, were pretreated with different amounts of IFN-ɑ2a, then infected with 17,000 FFU of CHIKV-mKate for 12 h; n = 7 separate wells infected on the same day; error bars represent sd. C-J. Cells were treated for 24 h by IFN, as indicated, followed by RT-qPCRs on ISGs. The data underlying this Figure can be found in **Supp Table 1**.

**Supplementary Figure 7.**
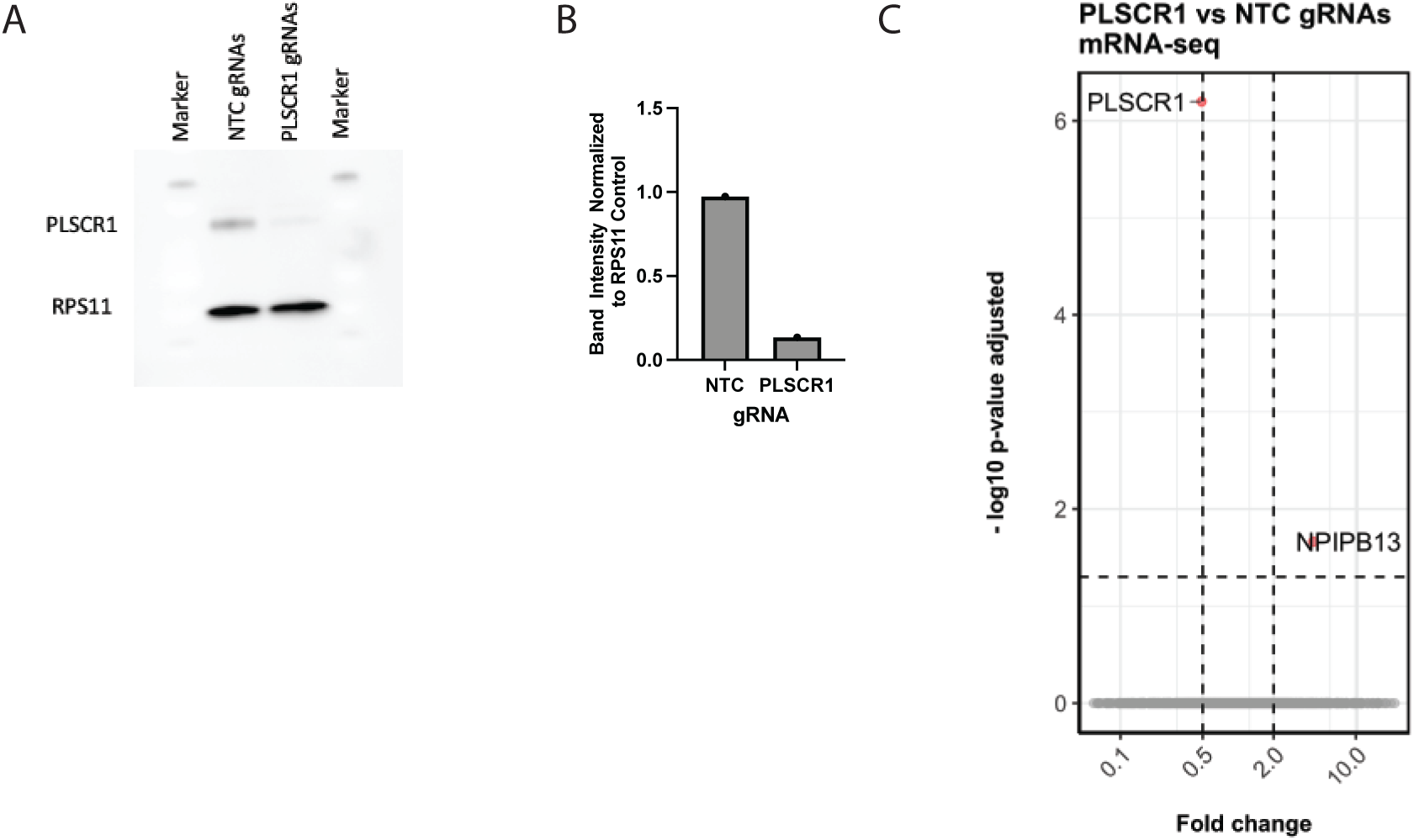
A. Western blot against PLSCR1 and RPS11. Cas-9-expressing Huh-7.5 cells were transfected with 4-gRNA pools targeting PLSCR1 or non-template control (as indicated) and cells were in culture for 7 d. B. Quantification of bands intensity in (A). C. mRNA-seq on PLSCR1 KO Huh-7.5 cells as in (A,B). The data underlying this Figure can be found in **Supp Table 1**.

**Supplementary Figure 8.**
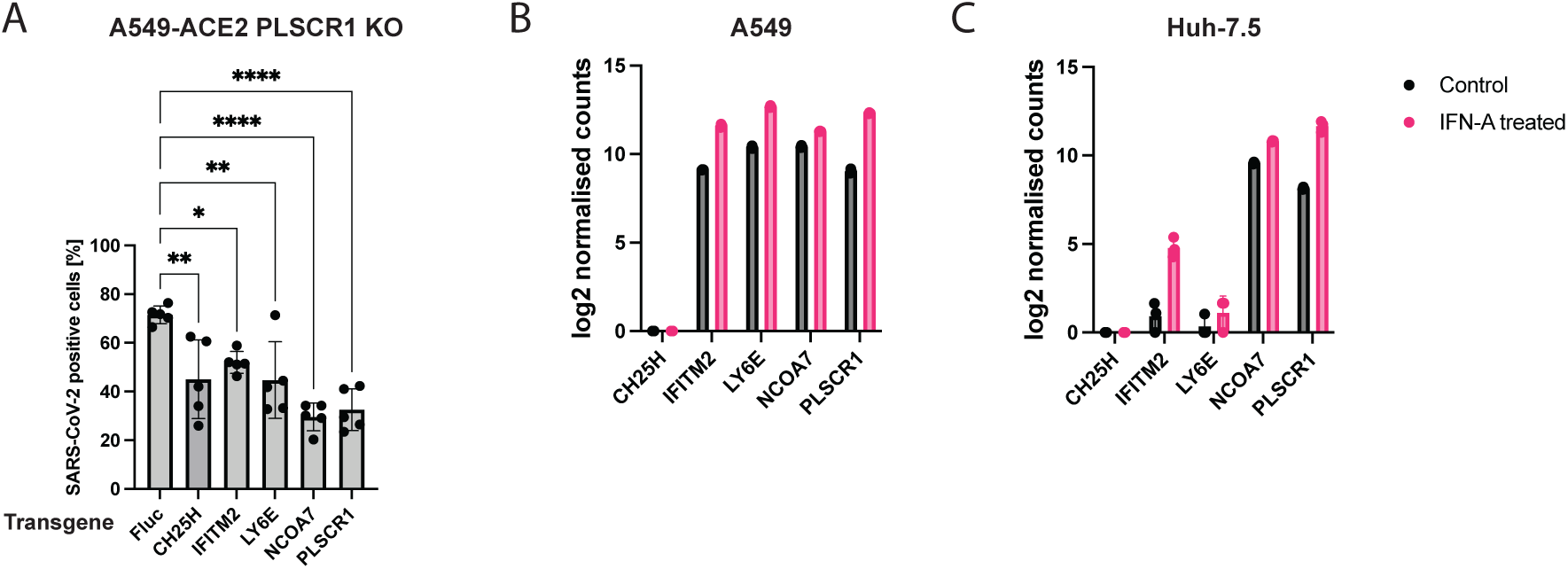
A. A549-ACE2 PLSCR1 KO cells were transduced to stably and ectopically express the indicated genes. The cells were then infected for 24 h with parental SARS-CoV-2. SARS-CoV-2 N was stained by IF and the percentage of positive cells determined by imaging. n = 4 separate wells infected on the same day. Error bars represent sd; **, p ≤ 0.01; ****, p < 0.0001; one-way ANOVA. B. Normalised reads count from mRNA-seq on A549 cells treated or not with 0.5 nM IFN-α2a for 24 h. C. Normalised reads count from mRNA-seq on Huh-7.5 cells treated or not with 0.5 nM IFN-α2a for 24 h. The data underlying this Figure can be found in **Supp Table 1**.

**Supplementary Figure 9.**
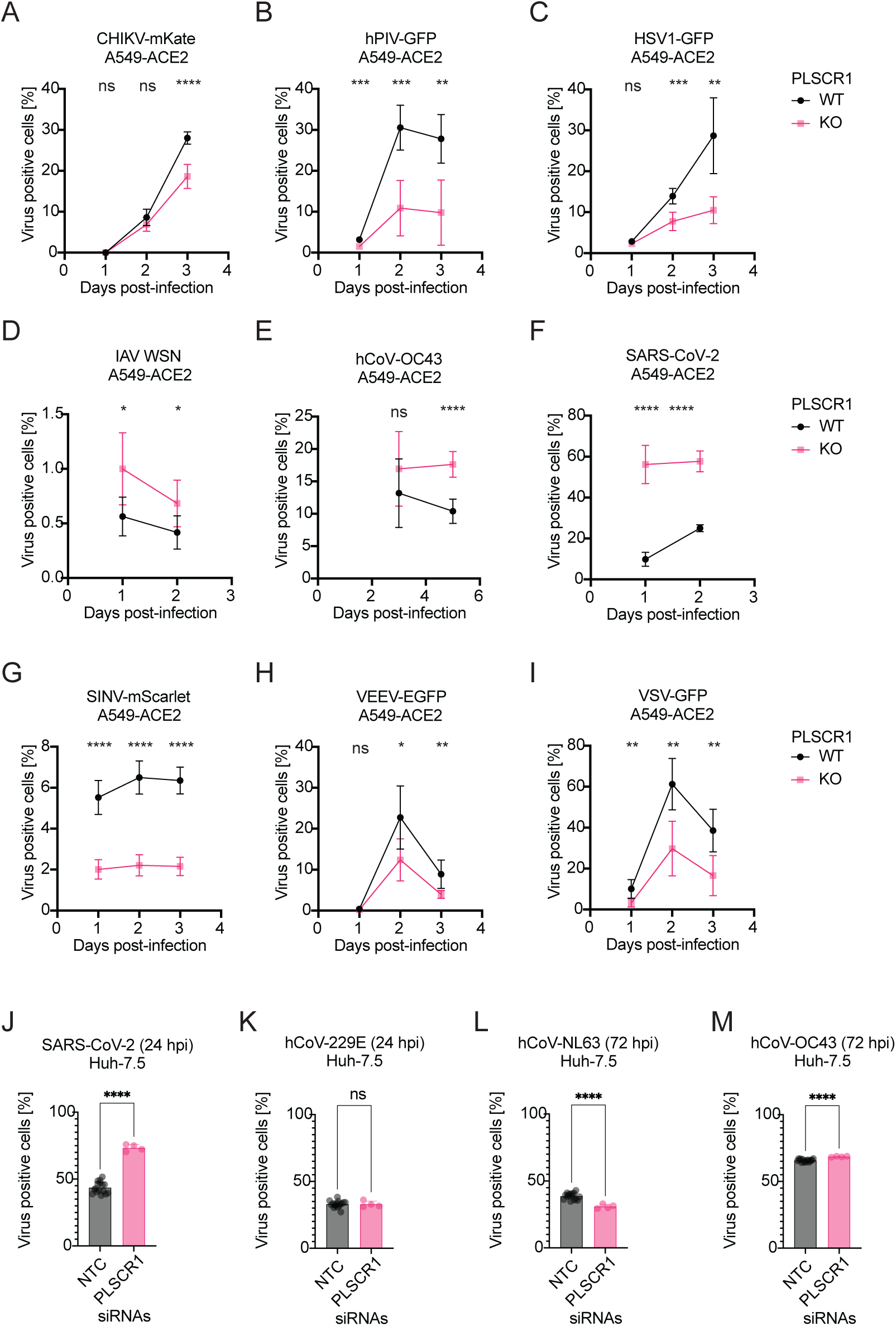
A-I. Infection of A549-ACE2 cells with viruses as indicated. IF staining was used for IAV WSN, hCoV-OC43 and SARS-CoV-2, otherwise a fluorescent reporter was used. Percentage of virus positive cells was determined by imaging. n = six separate wells infected on the same day, error bars represent SD; ns, non-significant; **, p ≤ 0.01; ***, p ≤ 0.001; ****, p ≤ 0.0001; ****, p ≤ 0.0001; two-tailed t-test. J-M. Infection of Huh-7.5 cells with viruses as indicated. siRNA knockdown of PLSCR1 vs non-template control (NTC). n = four to sixteen separate wells infected on the same day; ns, non-significant; ****, p ≤ 0.0001; two-tailed t-test. The data underlying this Figure can be found in **Supp Table 1**.

**Supplementary Figure 10.**
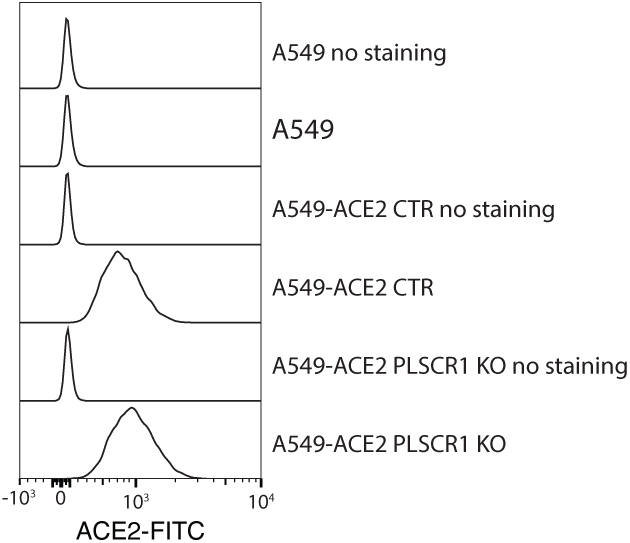
Flow cytometry measuring ACE2 surface levels in live A549-ACE2 cells.

**Supplementary Figure 11.**
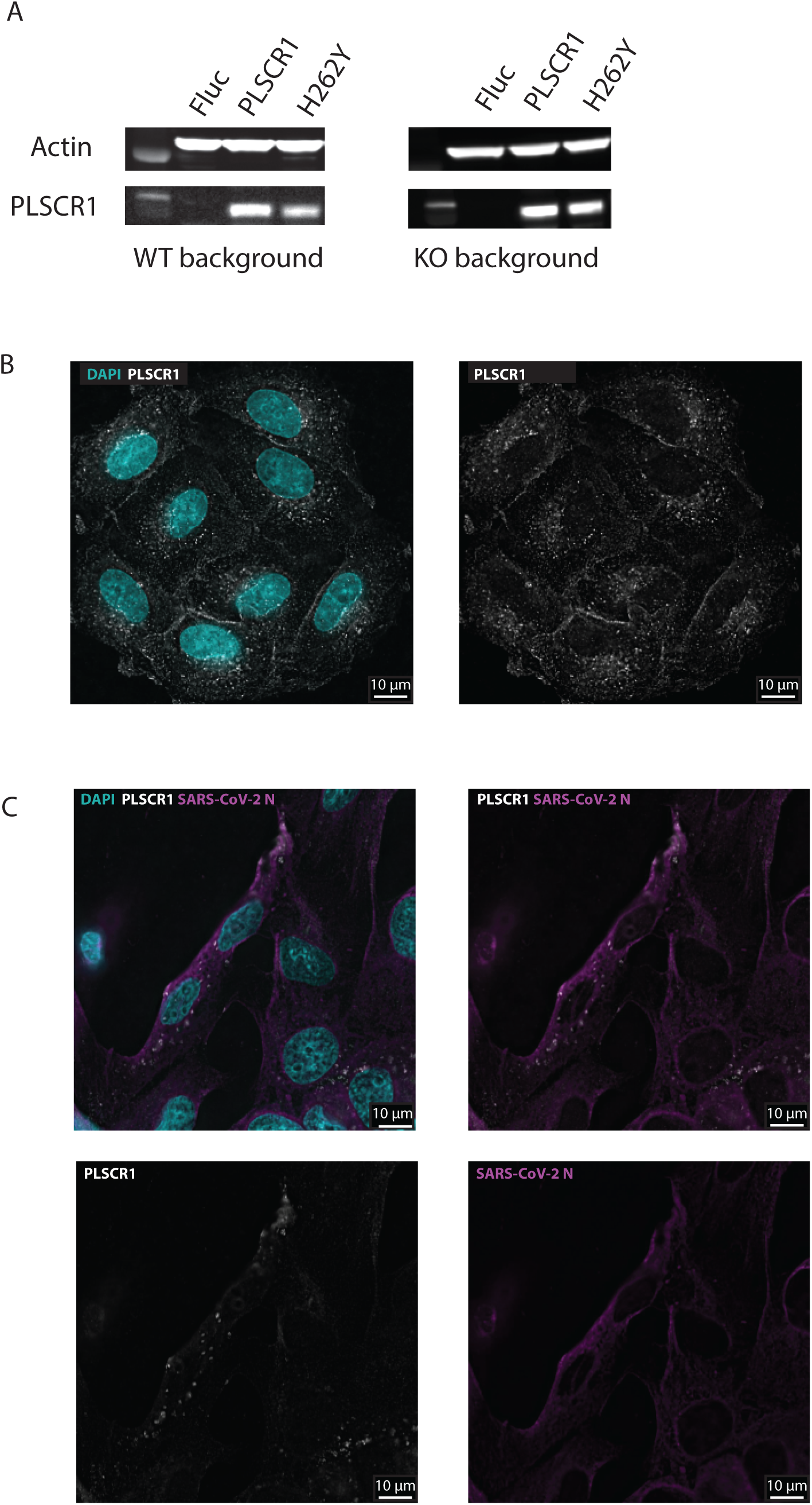
A. Western Blot against PLSCR1 and β-actin (loading control). A549-ACE2 PLSCR1 KO cells were stably transduced with FLAG-tagged Fluc, FLAG-tagged PLSCR1, or FLAG-tagged PLSCR1 H262Y. B. Representative confocal images of A549-ACE2 cells treated with IFN-α2a for 48 h. C. Representative confocal images of A549-ACE2 cells infected with SARS-CoV-2 for 24 h.

**Supplementary Figure 12.**
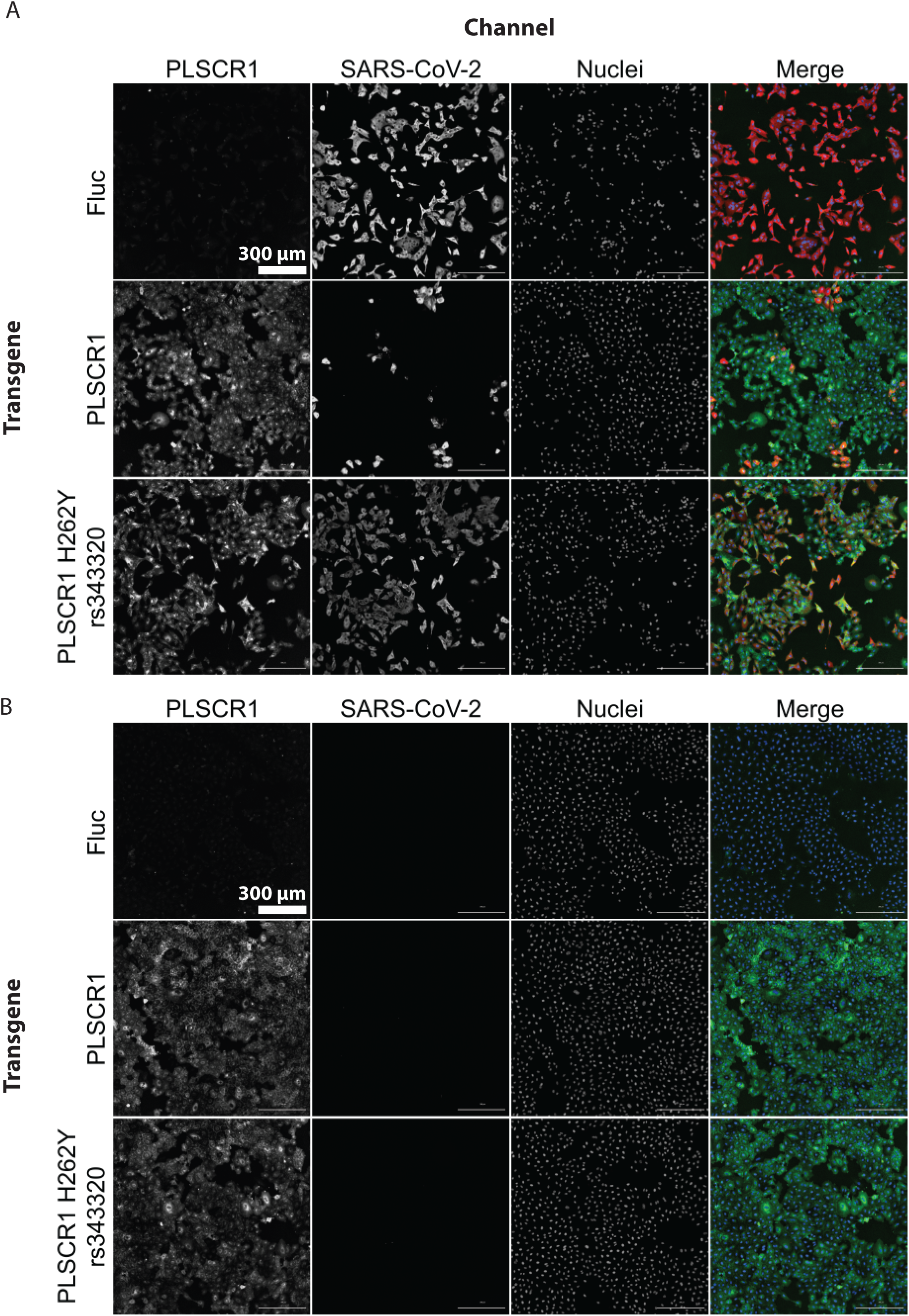
Representative images from the experiment shown in **Fig 7C**. A549-ACE2 PLSCR1 KO cells were transduced to stably and ectopically express the indicated PLSCR1 variants. The cells were then infected for 24 h with SARS-CoV-2 (A) or mock (B). SARS-CoV-2 N and PLSCR1 were stained and imaged

**Supplementary Figure 13.**
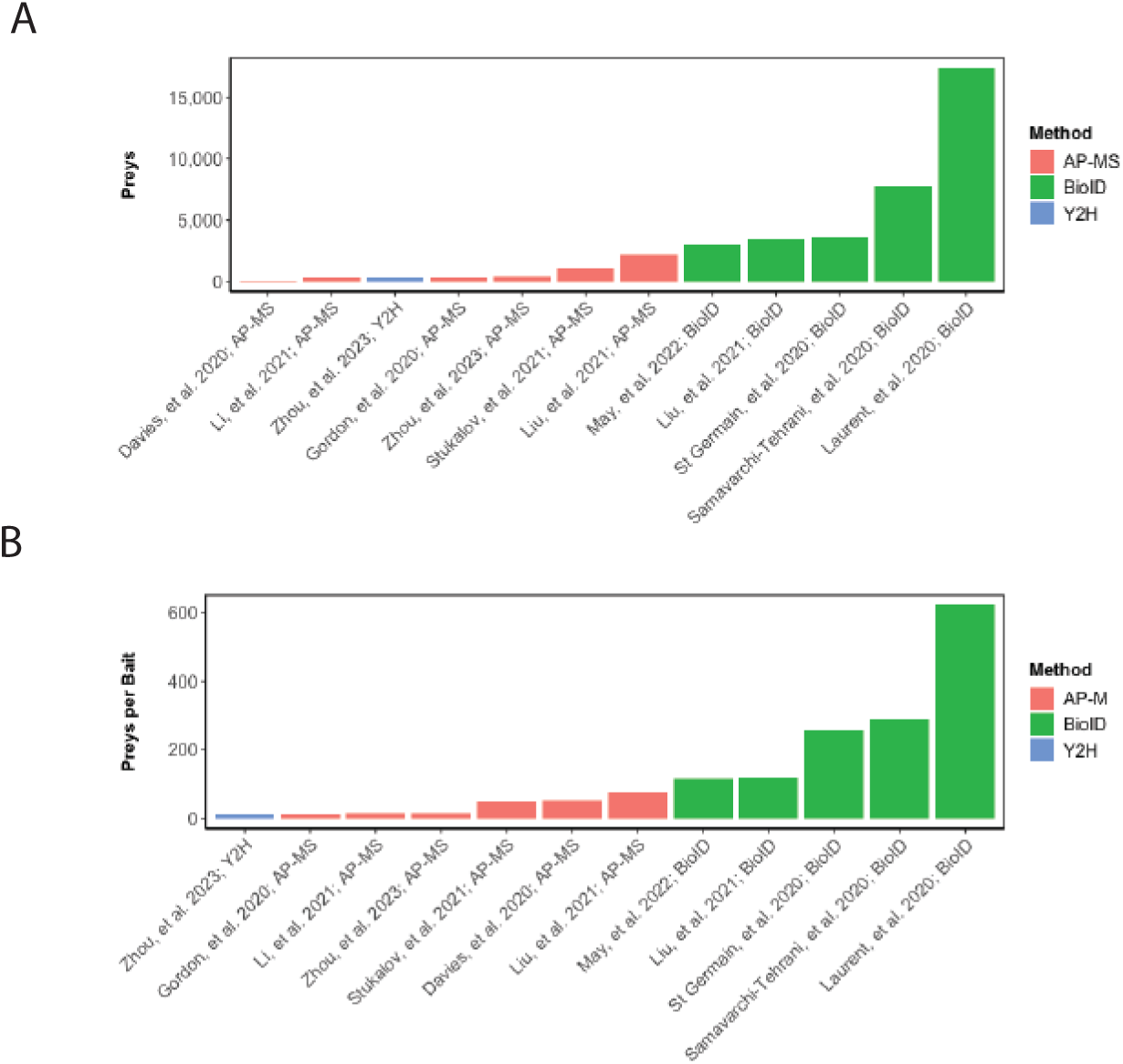
Comparison between selected SARS-CoV-2 protein interactome studies [100–102, 104–109, 157].

**Supplementary Figure 14.**
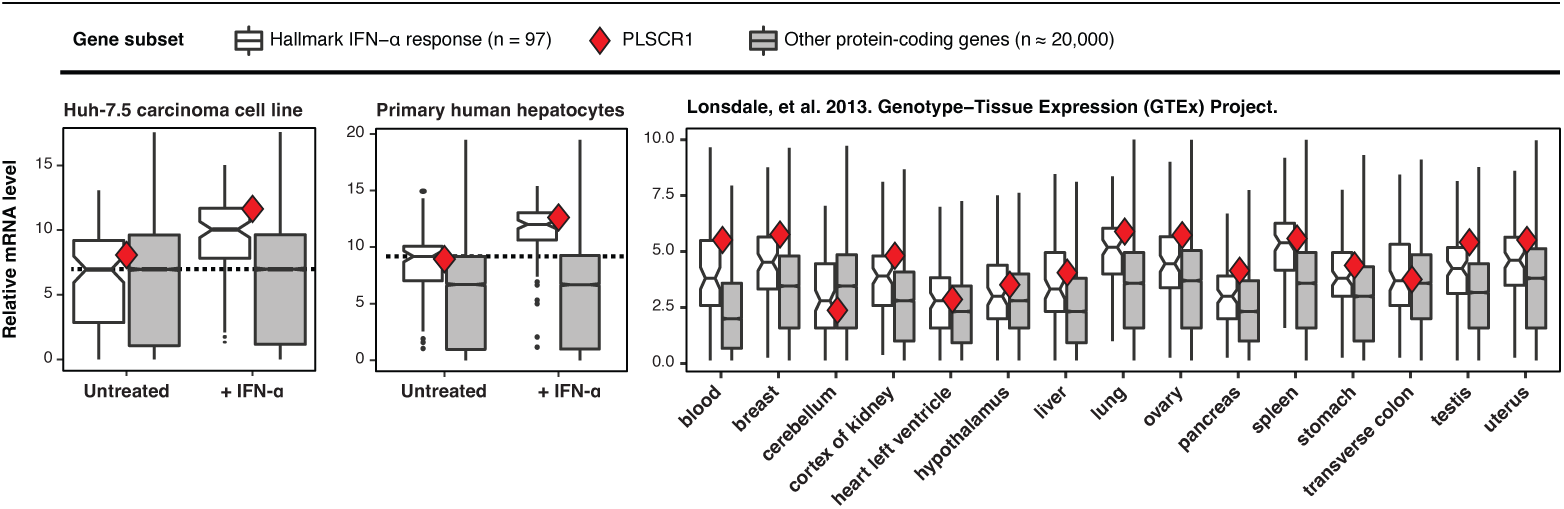
Comparison between the relative mRNA levels of 97 hallmark IFN-α-stimulated genes and the remaining transcriptome in cell lines as indicated. Red diamond, *PLSCR1* RNA level. For Huh-7.5 cells mRNA-seq, cells treated with 0.5 nM IFN-α2a for 24 h as in **Fig 2B**. For primary human hepatocytes, cells were treated with 0.1 nM IFN-α2a for 24 h (full data and methods to be released elsewhere). For the human tissues, data from [49]. The data underlying this Figure can be found in **Supp Table 1**.

## Supplementary Tables

Available on Dryad (DOI: 10.5061/dryad.6q573n65k).

**Supplementary Table 1.**

Data underlying Figures 1ABCEF, 2, 4ABC, 5CDEFGHIJKLM, 6ABCDEFHIJ, 7BCDE, S2ABC, S5BD, S6ABCDEFGHIJ, S7BC, S8ABC, S9ABCDEFGHIJKLM, S14

**Supplementary Table 2.**

mRNA-seq on SARS-CoV-2 infected Calu-3 and Huh-7.5 cells, Normalized reads, related to **Fig 1A**.

**Supplementary Table 3.**

mRNA-seq on SARS-CoV-2 infected Calu-3 and Huh-7.5 cells, Differential Gene Expression.

**Supplementary Table 4.**

mRNA-seq on IFN-treated Calu-3 and Huh-7.5 cells, Normalized reads, related to **Fig 1B**, and **Supp Figs 8C and 13**.

**Supplementary Table 5.**

mRNA-seq on IFN-treated Calu-3 and Huh-7.5 cells, Differential Gene Expression, related to **Fig 1F** and **Supp Fig 2C**.

**Supplementary Table 6.**

Genes expressed in three cell lines relevant for SARS-CoV-2 research (A549, Calu-3, Huh-7.5 cells) and human lung cells [47–49].

**Supplementary Table 7.**

Arrayed CRISPR KO screen, raw data.

**Supplementary Table 8.**

Arrayed CRISPR KO screen, analyzed data. Related to **Fig. 1F** and **Supp Fig 2B**. Columns are as follow: *Stat.W1_SARS2_IFN_0pM*: cell growth z-score determined by nuclear staining in samples with no interferon pretreatment and with SARS-CoV-2 infection. *Stat.W1_SARS2_IFN_1pM*: cell growth z-score determined by nuclear staining in samples with interferon pretreatment and with SARS-CoV-2 infection. *Stat.W1_Uninfected_IFN_0pM*: cell growth z-score determined by nuclear staining in samples with no interferon pretreatment and no SARS-CoV-2 infection. *Stat.W2_SARS2_IFN_0pM*: SARS-CoV-2 infection z-score in samples with no interferon pretreatment and with SARS-CoV-2 infection. *Stat.W2_SARS2_IFN_1pM*: SARS-CoV-2 infection z-score in samples with interferon pretreatment and with SARS-CoV-2 infection. *Stat.W2_Uninfected_IFN_0pM*: SARS-CoV-2 infection z-score in samples with no interferon pretreatment and no SARS-CoV-2 infection. Index: random index used for plotting. RNAseq: classification of the gene as an interferon-stimulated gene (“ISG”) or not (“NS”) based on the accompanying mRNA-seq data.

**Supplementary Table 9.**

Arrayed CRISPR KO screen, summary table.

**Supplementary Table 10.**

GSEA on arrayed CRISPR KO screen. Related to **Supp Fig 2D**.

**Supplementary Table 11.**

Consolidated list of human genes classified as hits in selected SARS-CoV-2 studies, full table. Related to **Fig 3** and **Supp Fig 3**.

**Supplementary Table 12.**

Consolidated list of human genes classified as hits in selected SARS-CoV-2 studies, summary table. Related to **Fig 3** and **Supp Fig 3**.

**Supplementary Table 13.**

Pathway analysis of hits from our arrayed screen alongside fifteen unbiased pooled screens [19, 31, 64–67, 69–71, 74, 75, 77–80] using the STRING database [208]. Related to **Supp Fig 4**.

**Supplementary Table 14.**

mRNA-seq on PLSCR1 KO Huh-7.5 cells, Normalized reads, related to **Supp Fig 7C**.

**Supplementary Table 15.**

mRNA-seq on PLSCR1 KO Huh-7.5 cells, Differential Gene Expression, related to **Supp Fig 7C**.

**Supplementary Table 16.**

mRNA-seq on IFN-treated A549 cells, Normalized reads, related to **Supp Fig 8B**.

**Supplementary Table 17.**

mRNA-seq on IFN-treated A549 cells, Differential Gene Expression, related to **Supp Fig 8B**.

**Supplementary Table 18.**

Plasmids used in this study.

**Supplementary Table 19.**

Gene fragments used in this study.

**Supplementary Table 20.**

Primers used in this study.

## Supporting Information

Available on Dryad (DOI: 10.5061/dryad.6q573n65k).

**Supporting Information 1**

Raw images, related to **Supp Figs 1B, 5AC**, **7A**, **11**.

**Supporting Information 2**

Flow cytometry gating strategy, related to **Supp Fig 10**.

**Supporting Information 3**

Flow cytometry raw data, related to **Supp Fig 10**.

**Supporting Information 4**

mRNA-seq raw reads, related to **Supp Tables 2, 3, 4, 5, 14, 15, 16, 17**.

## Notes

### Competing Interest Statement

The authors have declared no competing interest.

### Summary of Updates

We modified the title, abstract and author list. Code, minimally adjusted images, and flow cytometry data now available on Dryad (DOI: 10.5061/dryad.6q573n65k). RNA-seq data now available on NCBI (BioProject PRJNA1138251). We now supply the numerical values for all quantitative figures in Supp Table 1. We have clarified when the replicates from separate wells were infected on the same day. We discuss in more detail the number of essential genes and screen hits in the arrayed screen. We replaced kraken with Omicron (XBB 1.5) in the manuscript. We now present the screen control values separately in Fig 1E. Figure 2E (in the previous version) was moved to Supp Fig 2D (in the updated version). We created a new Fig 2 (in the updated version), that shows some pathways of interest in a more intuitive fashion (showing the z-score for each gene). We have clarified that the KO is PLSCR1 in the various panels of Figs 5 and 6 in the updated manuscript (Figs 6 and 7 in the previous version). Viral growth curves in Figs 6I and 6J in the updated version. We have included photos and rewritten the text related to Fig 6 in the updated version (Fig 7 in the previous version) for enhanced clarity. We have labeled PLSCR1 in Fig 7A in the updated version (Fig 8A in the previous version). Human genetics data updated in Fig 7. We describe the patient-derived fibroblasts in more detail in the results and methods section. We moved Figure 1 (in the previous version) to the supplemental section, Supp Figure 1A (in the updated version). We included the IFITM proteins. TMPRSS2 western blot (Supp Fig 1B). Single-cell RNA-seq data showing that PLSCR1 is constitutively expressed in SARS-CoV-2 target cells in vivo (Supp Fig 5D). Fig 5 (in the previous version) moved to Supp Fig 6 (in the updated version). Genetic association between PSCLR1 and some other ISGs blocking SARS-CoV-2 entry (Supp Fig 8). Added stats to A-I in Supp Fig 9 in the updated version (Supp Fig 5 in the previous version). PSCLR1 expression does not significantly affect ACE2 surface levels (Supp Fig 10). PLSCR1 H262Y does not alter the fact that PLSCR1 is primarily localized in the cytoplasm (Supp Fig 12). We discuss in more detail the different SARS-CoV-2 entry routes and implications for the PLSCR1 restriction. The Discussion section was modified to include the new data.

